# Ecological factors that drive microbial communities in culturally diverse fermented foods

**DOI:** 10.1101/2024.08.20.608727

**Authors:** Arya Gautam, Rahgavi Poopalarajah, Anique R. Ahmed, Bimarsh N. Rana, Tsedenia W. Denekew, Nakyeong Ahn, Lina Utenova, Yadu S. Kunwar, Nitin N. Bhandari, Aashish R. Jha

## Abstract

Fermented foods are increasingly recognized for their health benefits. Historically, cultures worldwide have relied on fermentation to preserve foods and enhance their digestibility, flavor, aromas, and taste. Despite the abundance of global diversity of fermented foods, the microbial communities in traditionally fermented non-European foods remain largely understudied. Here, we characterized the bacterial and fungal communities in 90 plant and animal based fermented foods from Nepal, South Korea, Ethiopia, and Kazakhstan, all traditionally prepared for household consumption. Our results reveal that these foods host diverse and intricately interconnected ecosystems of bacteria and fungi. Beyond the well-known fermenters such as lactic acid bacteria (LABs), Bacillales, and yeasts (Saccharomycetales), these foods contain additional microbes whose roles in fermentation are not well understood. While the microbial compositions of fermented foods vary by geography and preparation methods, the type of food substrate has the most significant effect on differentiating bacterial communities. Vegetable-based ferments harbor bacterial communities consisting primarily of LABs and potential pathways associated with carbohydrates degradation. Contrastingly, legumes and animal-based fermented foods are enriched with Bacillales and protein and lipid degradation pathways. Moreover, the microbial interactions, characterized via bacteria-bacteria and bacteria-fungi co-occurrence networks, differ significantly across traditionally fermented plants, legumes, and dairy products, indicating that microbial ecosystems vary between traditional fermented foods derived from different substrates. Our findings highlight the underexplored diversity of microbial communities in traditional fermented foods and underscore the need to understand the entire microbial communities present in these foods and their functions when evaluating their effect on nutrition and health.

## Introduction

Indigenous knowledge, previously overlooked and discredited, is now gaining recognition for its invaluable contributions to sustainability and stewardship [1]. Fermentation technology, originating from indigenous practices, has historically played a crucial role in sustainable food production and preservation. Over thousands of years, humans have utilized fermentation practices to improve food security, enhance digestibility, and augment taste [2]. Consumption of fermented foods has been linked not only to the evolution of the human brain [3] but it may also have contributed to the development of human taste receptors [4]. Recent studies have demonstrated that incorporating fermented food into our diet can reduce immune hyperactivity [5], promote gut health [6], and minimize chronic health conditions such as diabetes and hypertension [7]. These findings underscore the significant health benefits associated with the consumption of fermented foods.

Many of the fermented foods widely consumed today are the result of industrial fermentation processes, which heavily rely on a limited range of Lactic Acid Bacteria (LABs) and yeast strains that are known as starter cultures. The use of starter culture ensures consistent textures, flavors, and tastes in the final products [8]. In contrast, traditional fermentation methods often employ spontaneous fermentation by naturally occurring microbes or utilize a technique called “back-slopping,” where previously fermented products are used to initiate the fermentation process [9]. Consequently, traditional fermented foods are rich sources of LABs and yeasts that facilitate the fermentation processes and contribute to the maintenance of microbial community structures. For example, LABs such as *Levilactobacillus brevis*, *Lactiplantibacillus plantarum*, and *Lactococcus lactis* as well as yeasts including *Debaryomyces nepalensis* that are commonly found in traditional fermented foods produce proteins that inhibit the growth of undesirable microbes during food fermentation [10,11]. Moreover, several other bacteria and fungi present in traditional fermented foods may be involved in enhancing tastes, augmenting flavors, and improving nutrient quality. While LABs such as *Levilactobacillus brevis*, *Lactobacillus acetotolerans, Lactobacillus buchneri* and *Pediococcus pentosaceus* are known to produce sour flavors [12–16], members of the bacterial genera *Bacteroides* and *Pantoea* and fungi like *Kluyveromyces marxianus* generate aromatic flavor in traditional fermented foods [17,18]. Likewise, *Bifidobacteria* and non-pathogenic members of *Citrobacter* and *Klebsiella* synthesize vitamin B12 [19,20] while fungi such as *Cladosporium colombiae* and *Hannaella oryzae* produce the umami flavor in traditional ferments [21,22]. Despite their importance, many of the microbes in traditional fermented foods [17–43] remain markedly understudied. Therefore, it is crucial to comprehensively investigate the entire microbial communities present in traditional ferments to fully appreciate their relevance in food fermentation.

While there is ample literature describing the microbial communities found in industrially produced and artisanal fermented foods from Europe [44-48 and the references within], there remains a significant gap in our understanding of the microbes inhabiting fermented foods from non-European cultures. This includes their functions and the various factors influencing microbial community formation, such as preparation techniques and microbe-microbe interactions. In this study, we conducted 16S ribosomal RNA gene (16S rRNA) and Internal Transcribed Spacer 2 region (ITS2) amplicon sequencing to comprehensively characterize the microbial communities, assess their interactions, and explore their functional potential in 90 samples of 24 distinct fermented foods sourced from Nepal, South Korea, Ethiopia, and Kazakhstan that were prepared for household consumption. Our results reveal that traditionally fermented foods host a rich diversity of bacterial and fungal communities, which vary across the raw ingredients and preparation methods used. Co-occurrence network analyses demonstrated that the substrate contributes to both positive and negative associations between bacteria and fungi in these fermented foods. By integrating indigenous knowledge of food fermentation with genomics, our study provides a solid foundation for future efforts to characterize these microbes at finer taxonomic resolution, understand their functions, and optimize microbial communities to enhance safety, quality, aesthetics, and health benefits of fermented foods.

## Results

### Traditional fermented foods vary by raw ingredients and fermentation methods

We collected 90 samples of 24 different types of fermented foods from Nepal (n=64), South Korea (n=20), Ethiopia (n=5), and Kazakhstan (n=1) that were traditionally prepared for household consumption using diverse fermentation techniques **(Figure 1a, Supplementary Table 1**). These ferments encompass 53 vegetables, 13 legumes, 5 cereals, 14 dairy and 5 animal products (meat or seafood). The vegetable-based ferments include *achars* (n=24), *gundruk* (n=12)*, sinki* (n=2), and *taama* (n=4) from Nepal, *kimchi* (n=5), *oiji* (n=1), chilli paste (n=1), pickled plum (n=1), and plum juice (n=1) from South Korea, as well as *awaze* (n=1) and *datta* (n=1) from Ethiopia. The cereal based ferments included Ethiopian *injera* (n=1) and *difo dabo* (n=1) as well as the alcoholic beverage *chhyang* (n=1) and its amylolytic starter *marcha* (n=2) from Nepal. Fermented legumes consisted of Nepali *masyaura* (n=4), as well as fermented soybean pastes (n=5), and soy sauces (n=4) from South Korea. The dairy based ferments included 1 cheese sample from Kazakhstan, Ethiopian *ayib* (n=1), and Nepali *dahi* (n=4), *chhurpi* (n=3), and soft cheese (n=5) samples. We also included fermented meat products from Nepal called *sukuti* (n=3) and fermented seafood (n=2) from South Korea.

**Figure 1.**
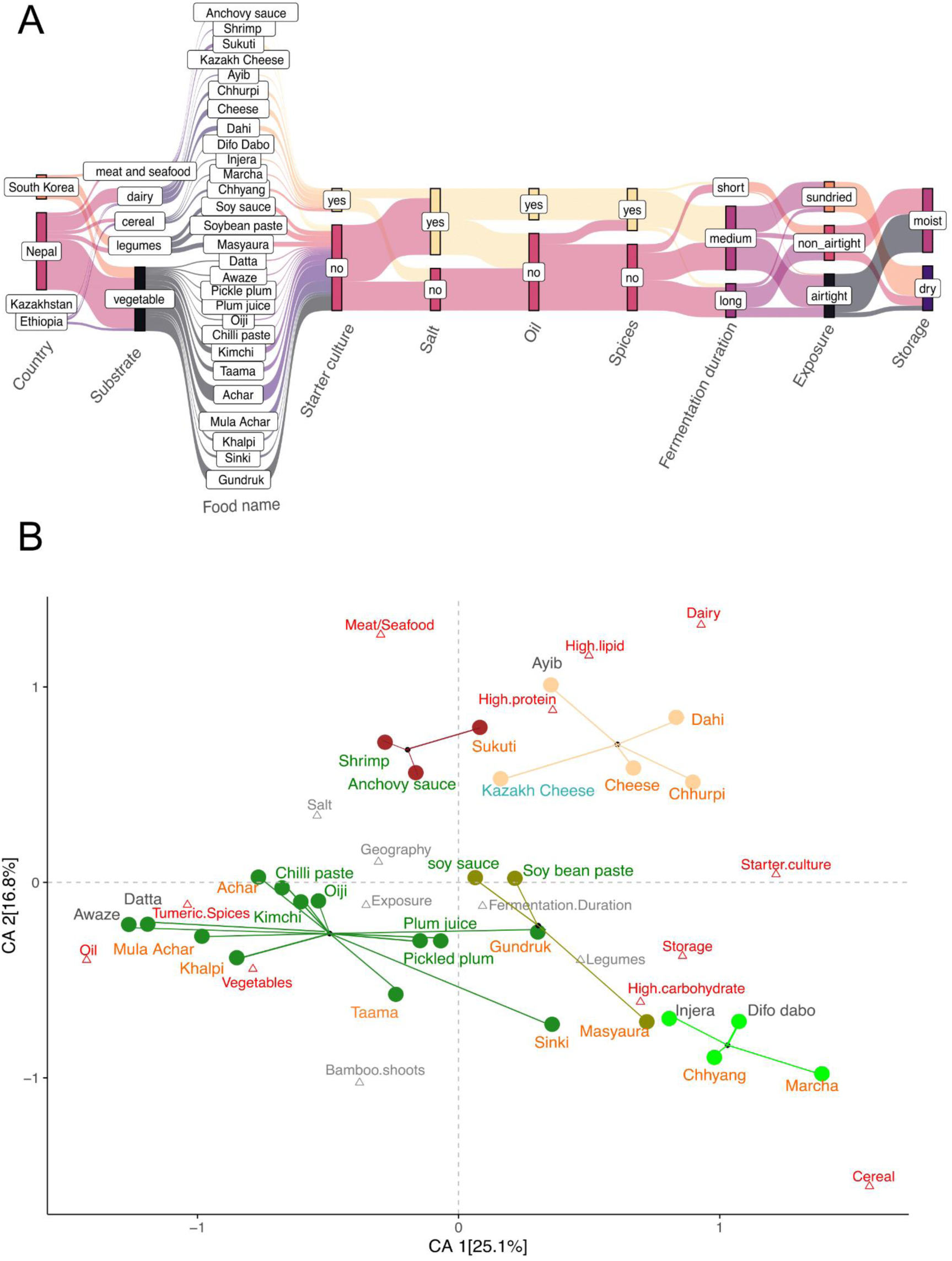
Diversity of raw ingredients and preparation methods used in traditional fermentation. We sampled 90 samples of 24 unique fermented foods from Nepal, South Korea, Ethiopia, and Kazakhstan. **(A)** Key parameters that vary during preparations of traditional fermented foods. Generally, Nepali *achars*, *khalpi, mula achar* and South Korean *kimchi, oiji,* and fermented chilli paste are prepared from garden vegetables marinated in turmeric, spices, oil, and salt that are spontaneously fermented by tightly packing and sealing in airtight containers. Nepali *gundruk* (mustard greens), *sinki* (radish), and *taama* (bamboo shoots) along with pickled plum and plum juice from South Korea, and *datta* and *awaze* from Ethiopia also rely on spontaneous fermentation. Legume based Nepali fermented foods such as *masyaura*, which are nuggets of ground black lentil paste mixed with taro, yam, or colocasia leaf and South Korean soy sauce and soybean paste are prepared without use of spices or oil. *Gundruk, sinki,* and *masyaura* are sun dried before storing at room temperature for up to one year. Fermented dairy products from Nepal, including *dahi* (yogurt), soft cheese, and *chhurpi* (dried and hardened cheese), Ethiopian *ayib*, and Khazakhi cheese are fermented using starter cultures. The cereal based fermented foods of Ethiopia included *injera* and difo dabo as well as the Nepali alcoholic beverage *chhyang*, which uses the amylolytic starter *marcha*, also rely on starter cultures. Fermented meat such as *sukuti* is marinated in spices and spontaneously fermented before sun drying. Similarly, South Korean fermented seafoods such as anchovy sauce and shrimp are fermented in salty brine solution. **(B)** Correspondence analysis of 17 abiotic variables associated with the different traditional fermented foods included in this study. Each food is represented by a circle colored by substrate: vegetables (dark green), dairy (orange), meat/seafood (maroon), cereal (lightgreen), and legumes (olive). The corresponding text labels are colored by the country of origin: Nepal (orange), South Korea (green), Ethiopia (grey) and Kazakhstan (blue). Each triangle represents an abiotic factor associated with food preparation and macronutrient profile. Factors in red contribute most to the top two dimensions in the CA. The first dimension of the correspondence analysis (CA1) primarily differentiates the spontaneously fermented vegetables and animal products mixed with oil, salt, and spices from foods fermented using starter cultures such as protein-rich dairy products (*dahi*, soft cheeses, and *chhurpi*) and cereal-based alcoholic beverage *chhyang* along with its starter *marcha*. Fermented plants (vegetables and cereals) differ from fermented animal-based products (meat, seafood, and dairy) along CA2.

Using current literature, we identified 17 variables describing macronutrient profiles and fermentation processes that distinguish the traditional fermented foods included in this study [49]. Multivariable analysis of these descriptors revealed that fermented foods from different regions of the world cluster together based on substrate type and preparation methods (**Figure 1b)**. Notably, plant-based foods such as fermented vegetables, legumes, and cereals exhibit significant differences from animal-based products. Within plant-based fermented foods, we observed differences between cereal-based, legume-based, and vegetable-based ferments. Similarly, there were variations between fermented dairy and fermented animal products (seafood and meat).

### Traditional fermented foods serve as natural reservoirs of diverse bacteria and fungi

To assess whether microorganisms in fermented foods permeate food surfaces, we aliquoted 64 samples of Nepali fermented foods into duplicate tubes before conducting DNA extractions. One aliquot of each fermented food sample underwent rigorous vortexing to dislodge surface microbiota, while the replicate samples were homogenized using a mortar and pestle to capture all microbes associated with the surface through the core of food samples. Microbial communities obtained from fermented food samples from both approaches differed significantly from the extraction controls (**Supplementary Figure 1**). But we observed no significant differences in the diversity, composition, and relative abundances of bacterial or fungal amplicon sequence variants (ASVs) between the homogenized and non-homogenized fermented food samples **(Supplementary Figure 1)**, indicating that microbes on the surface of fermented foods are indistinguishable from those present in the cores. Thus, we merged the sequencing reads obtained from homogenized and non-homogenized fermented food samples for subsequent analyses.

In total, we analyzed the bacterial communities in 90 fermented food samples using 16S rRNA sequencing. Additionally, we examined fungal diversity in 29 samples that yielded sufficiently high DNA concentrations for ITS2 sequencing (**Supplementary Figure 1**). Across the dataset, we identified a total of 104 bacterial and 91 fungal amplicon sequence variants (ASVs) exceeding relative abundance of 0.1%. Among the bacterial ASVs, 46 belonged to the order LABs, while 16 were members of the Bacillales order. Together, the LABs and Bacillales accounted for 78% of total bacterial reads (**Figure 2a)**. Likewise, Saccharomycetales emerged as the predominant fungal order in these fermented food samples, with less than half ASVs (40 of 91) comprising 71% of total fungal reads (**Figure 2b**). We classified these microbial taxa as “Canonical Fermenters” due to their well-established role in food fermentation [50–53]. On average, each sample harbored 35 (SD ± 4), 11 (SD ± 2), and 18 (SD ± 7) ASVs from LABs, Bacillales, and Saccharomycetales, respectively (**Supplementary Figure 2a**). While LABs, Bacillales, and Saccharomycetales were present in all samples, their relative abundances varied significantly within each food type (**Supplementary Figure 2b**, p=0.0002, 0.002, 0.004 respectively, *Brown-Forsythe test*). Fermented vegetables, dairy products, and alcoholic beverages were rich in LABs and Saccharomycetales, while fermented legumes, meats, and seafood showed elevated levels of Bacillales (**Figure 2c-d, Supplementary Figure 2b**). *Lactiplantibacillus sp*., a bacterial genera widely known for food fermentation [53], had the highest prevalence in our dataset and was found across all five substrate types (**Figure 2c**). *Pediococcus pentosaceus/Pediococcus stilesii,* known to produce sour flavor [12], was more prevalent in fermented vegetables and cereal, which are desirable qualities in these foods. *Issatchenkia orientalis*, known to tolerate highly acidic conditions and previously extracted from a variety of fermented foods [54,55], was the most prevalent fungi in our data. On the other hand, several members of yeasts from genera *Debaryomyces* dominated fermented dairy (**Figure 2d**).

**Figure 2.**
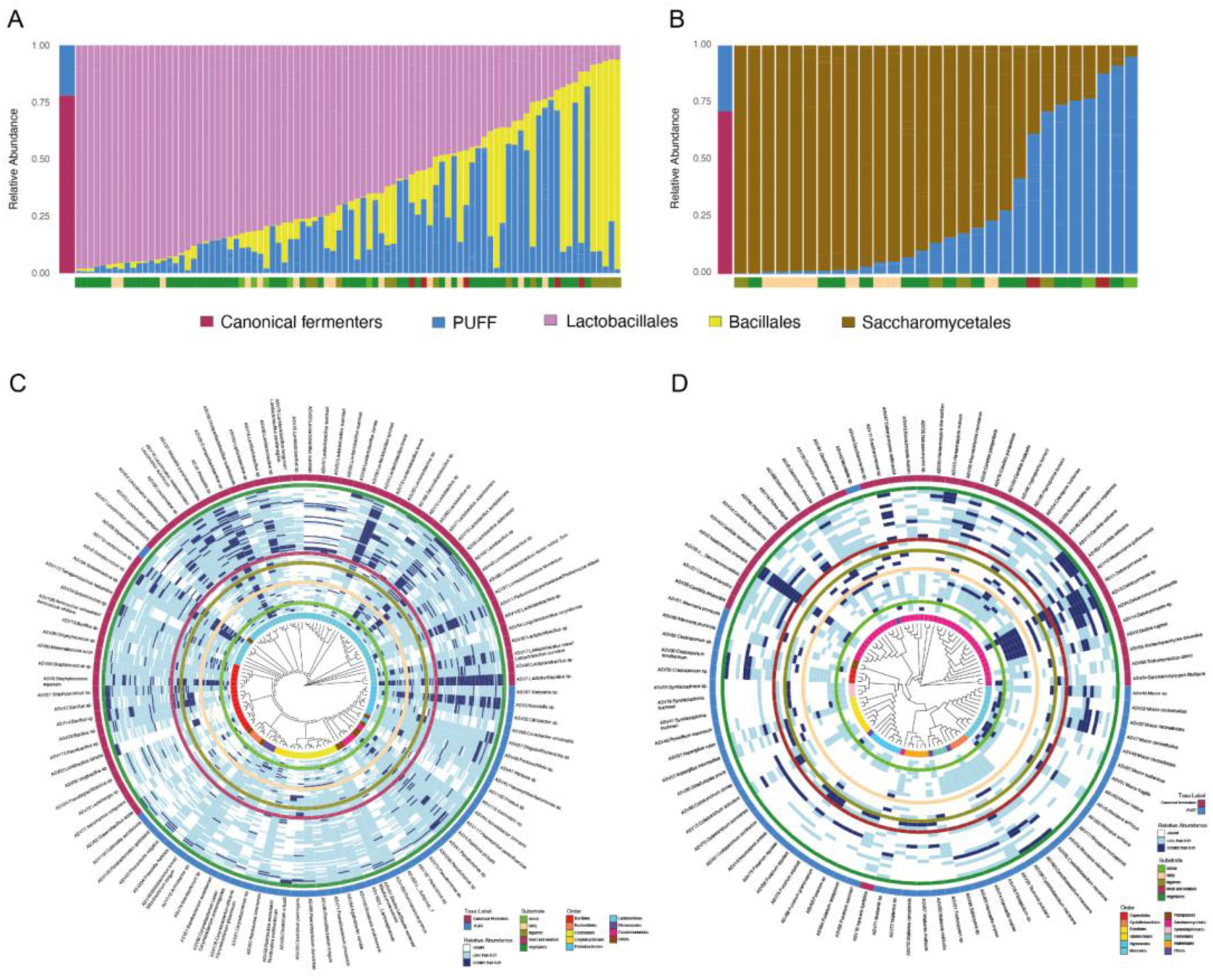
Traditional fermented foods exhibit heterogeneity in their microbial composition. **(A)** Leftmost column shows the relative abundances of Canonical Fermenters (Lactobacillales and Bacillales) across all samples in maroon and bacteria whose roles are Previously Undefined in Food Fermentation (PUFF) in blue. Subsequent columns show the relative abundances of Lactobacillales (pink), Bacillales (yellow), and PUFF bacteria (blue) in each sample. The bottom bar indicates substrate type: vegetables (darkgreen), cereals (light green), legumes (olive), dairy (beige), and animal products (brown). **(B)** Leftmost column shows the relative abundances of Canonical Fermenters (Saccharomycetales) across all samples in maroon and PUFF fungi in blue. Subsequent columns show their relative abundances in each sample. The bottom bar indicates substrate type colored as described above. **(C-D)** Phylogenetic tree of the bacterial and fungal ASVs and their relative abundances across all samples. The innermost ring represents the taxonomic Order. Subsequent rings represent individual samples grouped under the five colored rings corresponding to the respective substrate types. The outermost rings indicate whether a specific ASV is classified as Canonical Fermenter (maroon) or PUFF (blue).

In addition to the canonical fermenters, a notable portion of both bacterial (22%) and fungal (29%) reads represented microbes whose roles are <u>p</u>reviously <u>u</u>ndefined in food fermentation (PUFF, **Figures 2c-d**). The bacterial and fungal ASVs representing PUFF were distributed across our samples originating from broad geographic regions (**Figure 2c-d**), consistent with their detection in diverse traditional fermented foods in previous studies [17–43]. Among the notable bacteria in this category are members of the genera *Brevibacterium*, *Corynebacterium*, and *Salinivibrio. Brevibacterium* and *Corynebacterium* were found at higher relative abundances in our cheese samples and they are known to produce red or orange colors in smear-ripened cheeses [31]. *Salinivibrio*, previously isolated from fermented seafood and soy products [32,33], was most abundant in the fermented shrimp and soy sauce samples we obtained from South Korea (**Supplementary Figure 2c**). Prominent PUFF fungi include ethanol-producing *Rhizopus arrhizu*s and *Mucor indicus* [43]. These fungi were most prevalent in *chhyang*, the Nepali cereal based alcoholic beverage and its amylolytic starter, *marcha*. Likewise, *Tausonia pullulans,* primarily reported in *sauerkraut* and *kimchi* [34,35], exhibited highest relative abundances in our vegetable samples. The fungus *Fusarium oxysporum*, which thrives on free amino acids in fermented meat products [56], was most abundant in our fermented meat samples (**Supplementary** Fig 2d). Approximately 10% of bacterial and fungal taxa in our dataset belonged to plant endophytes, human skin commensals, and soil inhabitants. A key microbe in this category include *Pseudomonas*, a soil-dwelling bacterium and common plant endophyte [57], which was detected in all traditional fermented foods in this study.

### Abiotic factors shape the microbial compositions of traditional fermented foods

To identify factors contributing to the heterogeneity of microbial communities in diverse fermented foods, we conducted Principal Coordinate Analysis (PCoA) using weighted UniFrac dissimilarity distances of bacterial ASVs, followed by PERMANOVA with 10 variables associated with preparation methods (**Figure 3, Supplementary Figure 3**). When analyzing all 104 bacterial ASVs, we detected a significant association between geography and bacterial community structure (**Figure 3a**, p = 0.0001, *PERMANOVA*). However, repeating these analyses after removing the ASVs classified as PUFF resulted in marked reduction in the effect of geography on bacterial composition of these fermented foods (**Figure 3a**, p = 0.38, *PERMANOVA*). These results indicate that geographical differences in fermented foods emerge from bacteria whose contributions in fermentation processes are not fully understood.

**Figure 3.**
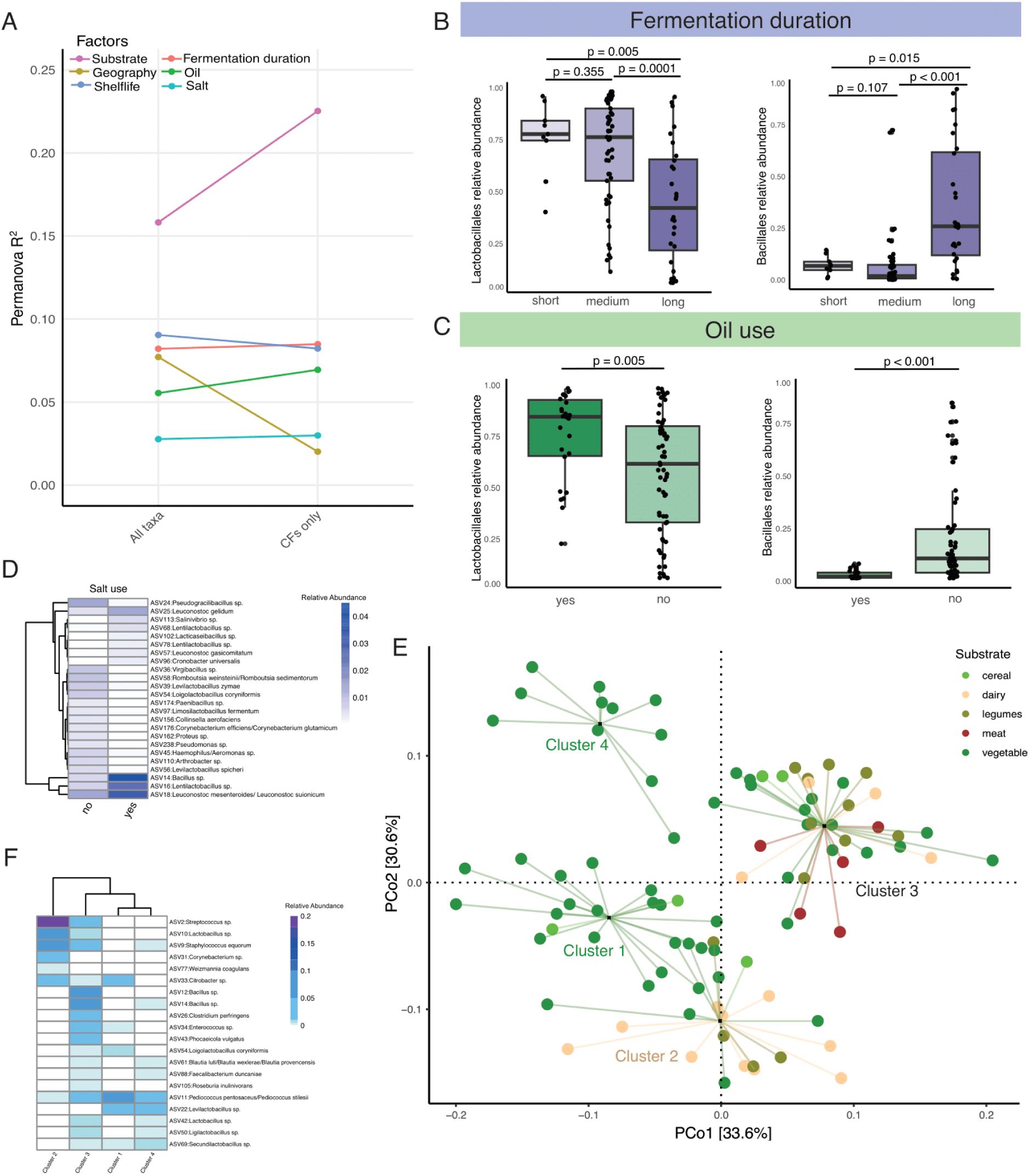
The bacterial compositions of traditional fermented foods are shaped by substrate type, preparation methods, and shelf life. **(A)** Permanova on the weighted UniFrac dissimilarity matrix identifies factors significantly associated with bacterial composition in traditional fermented foods. **(B)** The abundances of LABs decrease while that of Bacillales increase with longer fermentation periods. **(C)** Foods prepared without oil show an increase in the relative abundances of Bacillales and a decrease in LABs. **(D)** Differentially abundant ASVs between salted and unsalted fermented foods. **(E)** Principle Coordinate Analysis (PCoA) on the weighted unifrac dissimilarity matrix of traditional fermented food samples using 16S rRNA reads. Circles representing samples are grouped using PAM clustering and lines connect each sample to the cluster centroid. Colors represent substrate types: vegetables (darkgreen), cereals (light green), legumes (olive), dairy (beige), and animal products (brown).**(F)** Relative abundances of top 20 taxa with the highest variable importance factor (VIF) in the random forest model used to distinguish the four clusters of food samples.

Among the 10 variables included in the PERMANOVA, the substrate type exhibited the largest effect on the bacterial community structure of these fermented foods (**Figure 3a**, p<0.05, *PERMANOVA*). Unlike geography, the effect of food substrate on the bacterial composition of traditional fermented foods was more pronounced when the PUFF microbes were excluded. This indicates that the nutrients provided by the food substrates are the primary determinants of bacteria in traditional fermented foods.

In addition to substrate and geography, fermentation duration, the use of oil and salt, and shelf-life were also significantly associated with bacterial composition (**Figure 3a**, p<0.05, *PERMANOVA*). Foods fermented for longer periods and not immersed in oil constituted lower relative abundances of LABs and higher proportions of Bacillales (**Figure 3b-c**, p<0.05, *Mann Whitney test*). Similarly, we identified 24 bacterial ASVs that were differentially abundant between salted and unsalted foods (**Figure 3d**, FDR adjusted p<0.05 and absolute coefficient>2, MaAsLin2). Among these, *Salinivibrio sp.* and several members of *Bacillus*, *Lacticaseibacillus*, *Lentilactobacillus*, and *Leuconostoc* were more prevalent in salty foods. This finding is consistent with earlier studies that identified *Salinivibrio* and *Leuconostoc mesenteroides* as capable of thriving in high-salt environments [58,59].

Next, we performed partition-around-medoids (PAM) clustering using the first four axes of the bacterial principal coordinate analysis (PCoA), which revealed that the 90 samples of diverse fermented foods acquired from various geographical locations segregated into four broad clusters **(Figure 3e, Supplementary Figure 3)**. The first cluster predominantly comprised of spontaneously fermented fiber-rich vegetables and cereals from Nepal (p= 3.74e-05, *Fisher’s exact test*). The second cluster was enriched in dairy products (p=6.23e-05, *Fisher’s exact test*), while protein-rich legumes and animal products from Nepal and South Korea were overrepresented in the third cluster (p=3.82e-04, *Fisher’s exact test*). Notably, all fermented vegetables exceeding one year of shelf life were detected in the fourth cluster and showed higher relative abundances of bacteria associated with sour taste such as *Levilactobacillus brevis* and *Lactobacillus acetotolerans* [13–15], suggesting bacterial communities of fermented foods change over time. Notably, *Leviactobacillus brevis* is known to withstand low pH and has previously been reported in over ripened *kimchi* [15].

To validate these clusters independently, we employed a random forest classifier, which distinguished these four clusters with 100% accuracy (**Supplementary Figure 3)**. Furthermore, the bacteria with the highest variable importance factor (VIF) scores in the random forest classifier that were crucial for distinguishing the four clusters exhibited substrate specificity **(Figure 3f)**. For example, *Streptococcus sp., Corynebacterium sp., Brevibacterium sp*., and *Weizmannia coagulans*, previously associated with fermented dairy products [31,60,61], distinguished Cluster 2, which was enriched for the dairy samples. Likewise, *Bacillus sp.* prominent in soybean fermentation [62], delineated Cluster 3 comprising legumes and animal products. Finally, several LABs, known for their importance in vegetable fermentation [63], were prevalent in both Clusters 1 and 4 that were enriched for fermented vegetables. These results collectively indicate that despite differences in geographical origins and variations in the fermentation processes, substrate type has a significant contribution in shaping the bacterial composition of fermented foods.

When these analyses were conducted using the fungal ASVs, neither geography nor substrate type showed significant associations with the fungal compositions of these fermented foods **(Supplementary Figure 4**, p<0.05, *PERMANOVA*). Clustering analyses revealed these fermented foods can be grouped into three clusters, which was verified by a random forest classifier **(Figure 4a, Supplementary Figure 4)**, but these clusters were not linked to substrate types (p>0.05, *Fisher’s exact test*). The fungal species with the highest VIF scores **(Figure 4b)** also did not show substrate specificity **(Supplementary Figure 4)**. Instead, fermentation duration and the use of starter cultures appeared to be the major determinants of the fungal communities in these fermented foods **(Figure 4c)**. A dispersion analysis revealed that fungal communities in foods fermented using starter cultures showed greater uniformity relative to the spontaneously fermented foods (**Figure 4d**, p<0.05, *Mann Whitney test*). To determine whether the lack of association of substrate and geography with the fungal community structure was due to a smaller sample size of the fungal dataset, we analyzed the bacterial ASVs data from the same 29 samples. Despite the reduced sample size, bacterial composition remained significantly associated with substrates **(Supplementary Figure 5**), indicating that different abiotic factors contribute to bacterial and fungal diversity in traditional fermented foods.

**Figure 4.**
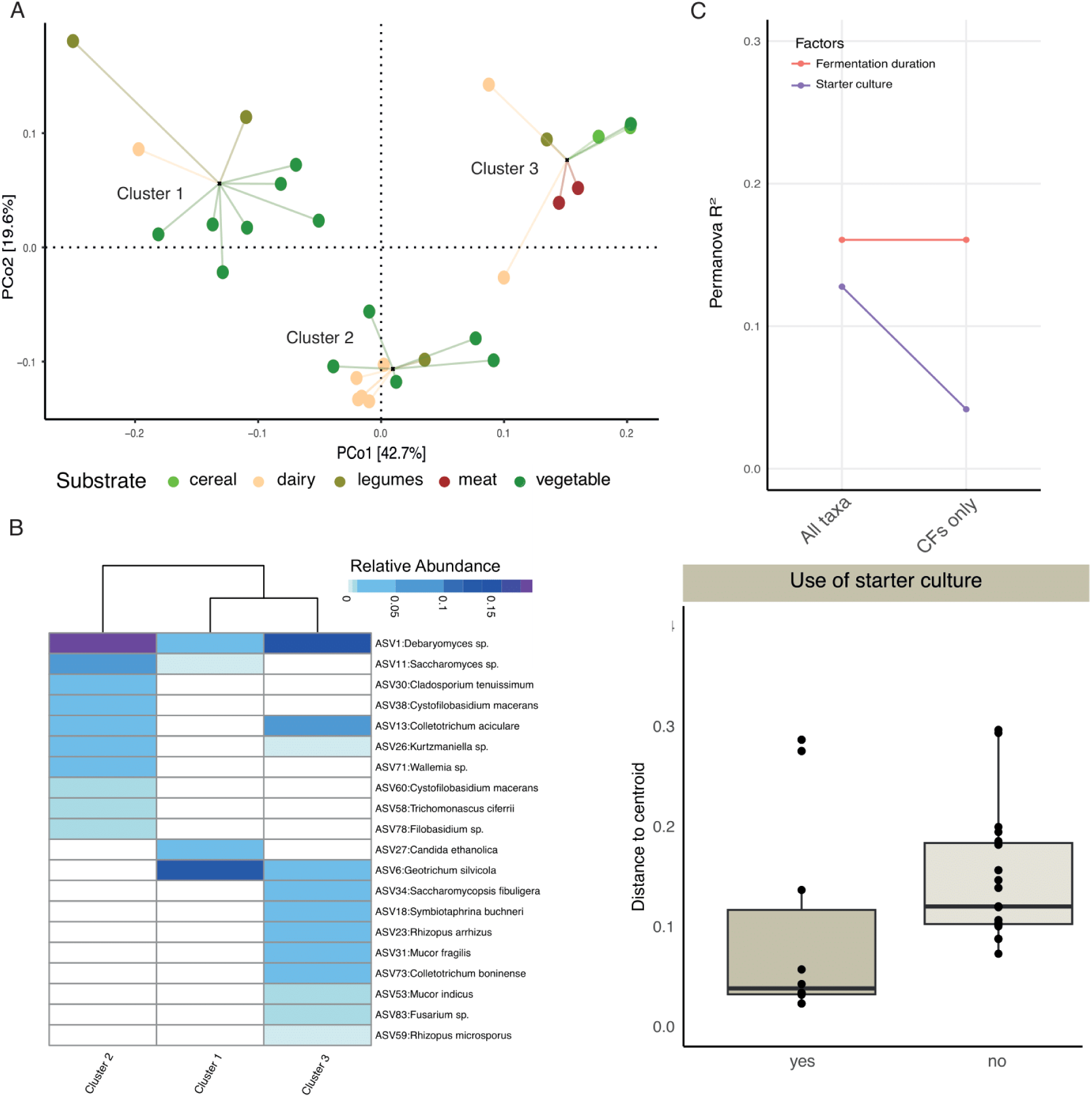
The fungal compositions of traditional fermented foods are shaped by fermentation duration and use of starter cultures. **(A)** Principle Coordinate Analysis (PCoA) of the weighted UniFrac dissimilarity matrix of traditional fermented foods using ITS2 reads followed by PAM clustering. Samples are represented by circles color coded by substrates and linked to the cluster centroids by lines. **(B)** Relative abundances of top 20 taxa with the highest VIF in the random forest model used to distinguish three food clusters. **(C)** Permanova on the weighted UniFrac dissimilarity matrix identified factors significantly associated with the fungal composition. **(D)** Dispersion analysis using weighted UniFrac distances comparing spontaneously fermented foods with those fermented using starter cultures.

### Bacterial pathways differ by fermented food substrates

Considering the significant role of substrate in shaping bacterial communities in traditional fermented foods, we sought to examine whether the bacterial communities in different substrates also differ functionally. Using Phylogenetic Investigation of Communities by Reconstruction of Unobserved States (PICRUSt2) [64], which enables prediction of functional content of microbial communities from amplicon sequencing data, we inferred the relative abundances of Kyoto Encyclopedia of Genes and Genomes (KEGG) [65] pathways in our fermented foods. Comparison of KEGG pathway relative abundances revealed distinct functional profiles across different substrates (FDR adjusted p <0.05, LinDA, **Figure 5, Supplementary Table 3,4**). Bacteria in vegetable- and cereal-based fermented foods were enriched for several pathways involved in carbohydrate metabolism (PPP), folate biosynthesis, as well as retinol (Vitamin A) and thiamine (Vitamin B1) metabolism. Plants contain high fibers, which provides the ideal substrate for bacteria that metabolize carbohydrates. Folate, an essential micronutrient necessary for cell growth and red blood cell formation [66], is often recommended during pregnancy to reduce the risk of neural tube defects in infants [67]. Retinol is important for vision, cellular development, and immunity [68], while thiamine is vital for glucose metabolism, as well as nerve, muscle and heart function [69]. Similarly, compared to plant based fermented foods, bacterial communities in fermented dairy, legumes, and animal products showed higher abundances of pathways involved in amino acid metabolism consistent with their macronutrient profiles. They were also enriched for Vitamin B6 metabolism, which is vital for nervous and immune system functions [70]. These findings highlight that microbes present in different fermented foods possess metabolic capabilities that match macronutrient profiles of substrates and may contribute biosynthetic compounds that are relevant to human health.

**Figure 5.**
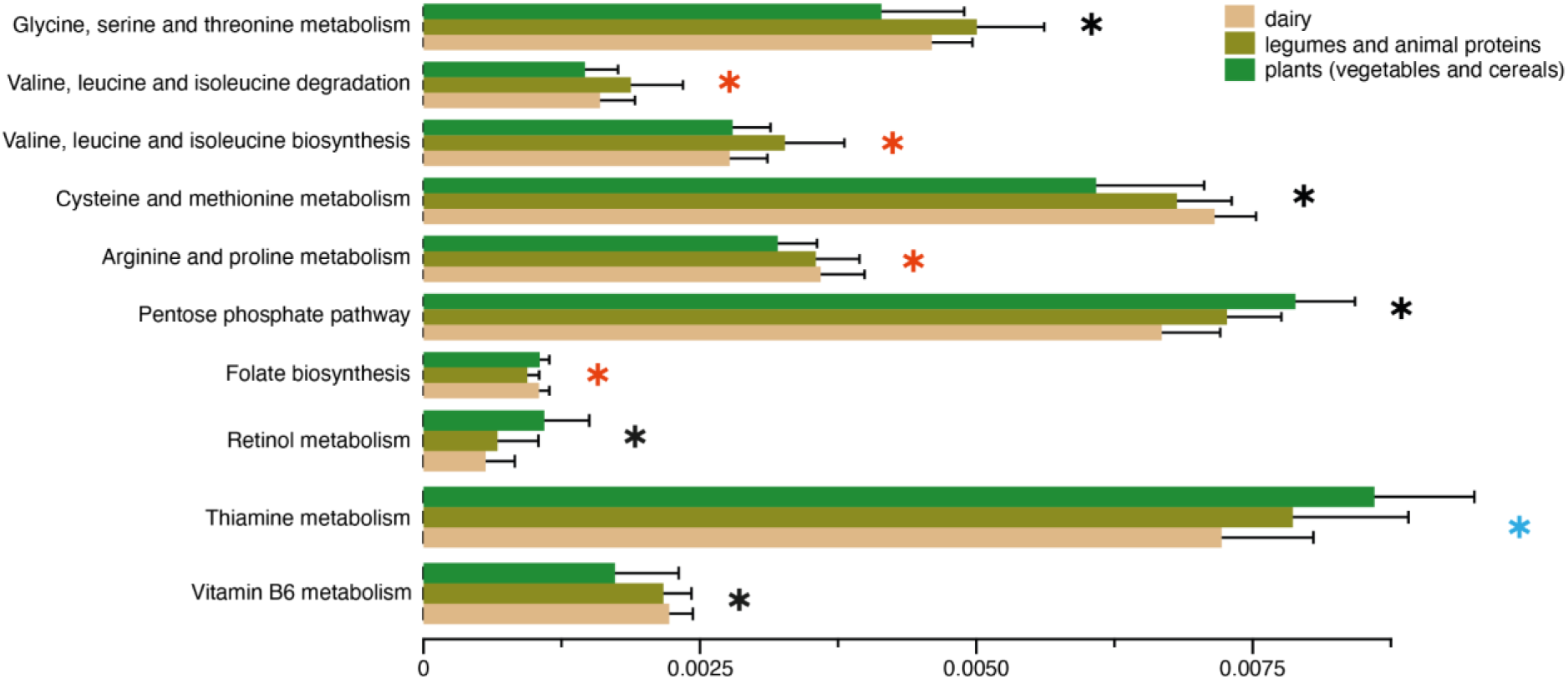
Bacterial functions may differ across food substrates. Relative abundances of putative functional pathways that are significantly different among substrate types (LinDA, p adjusted <0.05). Pathways with a blue star are significantly different between plants and dairy, those with a red star are significantly different between plants versus dairy, legumes and animal products, and those with a black star are significantly different across all three substrate types.

### Biotic interactions influence fermented food microbial communities

Microbial interactions play a pivotal role in shaping the microbial communities in fermented foods, consequently influencing their community structure and functions. Some microbes act as primary decomposers, breaking down food substrates to supply nutrients and metabolites to neighboring microbes, thereby facilitating their growth [71]. Others produce toxins that inhibit the proliferation of nearby microbial cells, shaping the overall community composition [72]. To elucidate the microbial interactions in traditional fermented foods, we constructed co-occurrence networks using 104 bacterial ASVs from all 90 food samples and 91 fungal ASVs from the 29 ferments in our dataset. Our analyses revealed densely interconnected bacteria-bacteria and fungi-fungi networks within these fermented foods **(Supplementary Figure 6)**. We identified five distinct co-abundance groups (CAGs) for both bacteria and fungi that varied in connectedness and transitivity. Connectedness measures the number of edges (co-occurrences) of each node (ASV) within a CAG, normalized by the total number of nodes in the CAG. Transitivity assesses how often two interconnected nodes co-occur with a common node. **(Supplementary Figure 7).** Both metrics evaluate how densely ASVs are interconnected within each CAG. We also identified focal microbes within each CAG by assessing their centrality using three measures: strength, betweenness, and eigenvector centrality **(Supplementary Figure 8)**. Microbes with high centrality are indicative of keystone species in microbial ecosystems [73].

Among the bacterial CAGs, the *Lactobacillus delbrueckii*-CAG emerged as the most interconnected and showed highest transitivity. It consisted of *Lactobacillus delbrueckii* and two PUFFs, *Brevibacterium sp.* and *Corynebacterium sp.* as focal members of this CAG. The *L. delbrueckii*-CAG was predominantly found in dairy products within our dataset **(Figure 6a),** consistent with association of these bacteria with cheesemaking processes [31, 74]. A second bacterial CAG that was prominent in legumes, meat, and seafood in our dataset comprised members of *Bacillus* as the focal taxa along with *Tetragenococcus halophilus*, *Oceanobacillus sojae*, *Enterococcus sp*., and *Salinivibrio sp*, which is consistent with the roles of these bacteria in soy and protein fermentation [33,52]. In addition, several human gut associated bacteria that were categorized as PUFF, including *Faecalibacterium*, *Blautia*, and *Ruminococcus gnavus*, showed high centrality measures in this *Bacillus*-CAG. Conversely, the fermented cereals and vegetables in our dataset were distinguished by two bacterial CAGs that demonstrated high connectedness. One of them comprised *Loigolactobacillus coryniformis* and *Levilactobacillus suantsaii* as the focal species but it also consisted of other canonical fermenters such as *Leuconostoc mesenteroides/suionicum*, *Lactiplantibacullis sp*. as well as PUFF bacteria such as *Arthrobacter*, *Proteus* and human gut microbiome associated bacteria such as *Prevotella histicola* and *Bifidobacterium*. The second CAG, consisted of *Lactobacillus acetotolerans* as a focal species. *L. acetotolerans* is associated with sour taste in vinegar, pickles, and alcoholic beverages [13,75] and these are the desired traits in the fermented vegetables from Nepal and South Korea. The fermented vegetables with longer shelf lives had a markedly higher proportion of this CAG. Finally, the fifth bacterial CAG consisted of several bacteria commonly associated with plant endophytes and soil such as *Acinetobacter johnsonii*, *Ralstonia picketti,* and *Pseudomonas sp.* as central members. This CAG did not show substrate specificity and was present at low proportions across all substrates.

**Figure 6.**
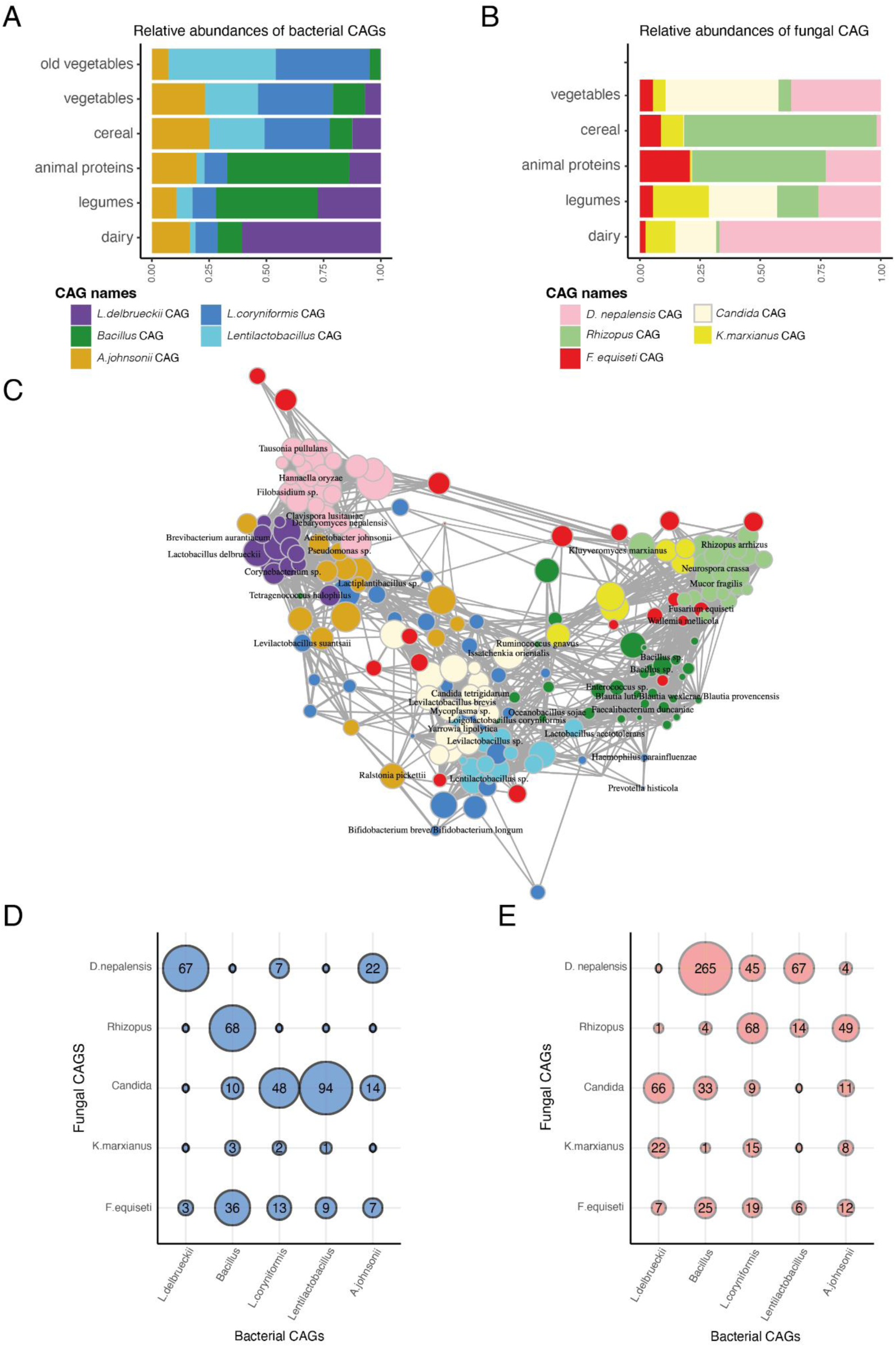
Co-occurrence analysis reveals densely interconnected microbial ecosystems in traditional fermented foods. (A,B) Relative abundances of bacterial and fungal co-abundant groups (CAGs) across substrate types. **(C)** Bacteria and fungi that significantly co-occur with one another in traditional foods. **(D,E)** The number of significant positive and negative associations between bacterial and fungal CAGs.

In addition to the positive interactions between bacteria within each CAG, we found antagonism between the five bacterial CAGs. The *L. delbrueckii*-CAG that defined the dairy bacterial community was highly antagonistic to the *Bacillus*-CAG prominent in legumes, meat, and seafood. Similarly, bacterial members of the two CAGs consisting of LABs and prevalent in vegetables showed several negative interactions with both the *L. delbrueckii*-CAG and the *Bacillus*-CAG.

Unlike bacteria, most of the fungal CAGs displayed a broader substrate distribution, with three prominent CAGs consisting of *Debaryomyces nepalensis*, *Candida*, and *Rhizopus* as focal species, respectively (**Figure 6b**). These CAGs demonstrated high connectivity and transitivity and were characterized by distinct focal species from the PUFF category that potentially play key roles within the microbial network. Some key focal members of the *D. nepalensis*-CAG consisted of several yeasts including *Tausonia pullulans*, *Clavispora lusitaniae*, *Filobasidium sp*., and the umami-associated *Hannaella oryzae* [21]. The *Candida*- CAG consisted exclusively of yeasts including low pH tolerant *Yarrowia lipolytica* and *Issatchenkia orientalis* that are associated with appearance, flavor, and taste of diverse fermented foods [54,55,76]. The *Rhizopus*-CAG included molds such as *Neurospora crassa* known to degrade allergens in soybean [77] and *Mucor fragilis* known for producing bioactive compounds [78]. The fourth CAG consisted of flavor-enhancing fermentation capable yeast, *Kluyveromyces marxianus* as a central species [18,79]. The final fungal CAG consisted of several environmental fungi including *Fusarium equiseti* and *Wallemia mellicola*. The *D. nepalensis*-CAG was highest in dairy (∼67%) but also detected at high proportions in vegetables (37%). The *Rhizopus*-CAG was high in fermented cereals and meat samples, while the *Candida*-CAG was detected at appreciable proportions in vegetables, legumes, and dairy. The remaining two fungal CAGs were distributed across the different substrates at low proportions.

Moreover, joint co-occurrence network analysis of bacterial and fungal ASVs revealed extensive symbiotic and antagonistic relationships across the two kingdoms, while maintaining the within-kingdom network architecture described above (**Figure 6c-e**). Notably, bacterial members of the *L. delbrueckii*-CAG that was highly prevalent in dairy frequently co-occurred with the fungi in the *D. nepalensis*-CAG and showed negative interactions with the *Candida*-CAG. In contrast, the *Bacillus*-CAG was highly antagonistic to *D. nepalensis*-CAG but co-occurred with members of *Rhizopus*-CAG. Furthermore, members of the two LAB dominated CAGs associated with fermented vegetables coexisted with fungi in the *Candida*-CAGs, which consisted of several acid tolerant species, indicating that both bacteria and fungi may contribute to create a low pH environment where specific bacteria and fungi can proliferate. Overall, our findings highlight the complex dynamics of microbes within fermented food communities and underscore their intricate interdependence that shape the microbial ecosystems in traditional fermented foods.

## Discussion

Fermenting foods is an age-old technique of immense cultural significance worldwide [2,80]. Various cultures worldwide have developed unique approaches for fermenting locally sourced raw ingredients [8,9]. These methods— honed through empirical knowledge passed down for generations –have yielded a myriad of fermented foods. Consequently, fermented foods have become integral to their culinary heritage [2]. These traditional fermentation techniques have yielded a diverse array of fermented foods constituting a range of substrates such as cereals, legumes, vegetables, dairy, meat, and seafood [9]. Traditional fermentation methods often involve soaking or boiling the raw ingredients, spontaneous fermentation or back-slopping, followed by sun-drying the ferments to extend the shelf life of food. In contrast, industrial fermentation often involves starter cultures. Understanding how different fermentation methods contribute to the microbial diversity of traditional fermented foods can reveal microbes that influence appearance, taste, flavor, and health benefits of these products. However, factors driving microbiome community formation in non-European fermented foods remain poorly understood. While microbial ecosystems of European foods have been extensively characterized using high-throughput genomics [44–48,81], comprehensive characterization of microbial identities in non-European fermented foods has been hindered due to the use of culture-based approaches [82–84]. Although recent studies have implemented metagenomics to assess microbial communities in traditional fermented foods [85–88], these works are often limited by small sample sizes and a lack of diversity of fermented food samples. Thus, there remains much to explore regarding the microbial ecosystems of non-European traditional fermented foods.

In this study, we sought to elucidate the ecological factors governing the microbial communities in traditional fermented foods commonly consumed in Asia and Africa that are derived from a broad range of food substrates and prepared using diverse fermentation techniques. Our results reveal several important aspects of fermented food microbiomes that are crucial for future studies. First, we used two different DNA extraction methods to demonstrate that both bacteria and fungi residing on food surfaces do not differ significantly from those present in the cores, which obviates the need for sample homogenization in future metagenomics studies. Homogenizing fermented foods would release large quantities of plant and animal DNA which can be a major detraction in whole metagenomics sequencing [45,89]. Second, the use of 16S rRNA and ITS2 amplicon sequencing allowed us to catalog the comprehensive diversity of bacteria and fungi present in the traditional fermented foods that extend beyond the Canonical Fermenters such as LABs, Bacillales, and yeasts that have well known functions in food fermentation. Third, our results indicate that the PUFF microbes, whose roles in food fermentation remain relatively unexplored, are important contributors in food fermentation. For example, the PUFF microbes and not the Canonical Fermenters differentiated the fermented foods originating from diverse geographic regions, suggesting that these microbes may be the primary contributors to regional differences in flavor and taste in fermented foods. Several of them were identified as focal members in the bacterial and fungal CAGs, indicating they may be keystone species in microbial ecosystems of traditional ferments. Moreover, enrichment of specific PUFF microbes in fermented foods from broad geographic distributions but sharing specific preparation methods, such as *Salinivibrio* in salted foods, *Brevibacterium* and *Corynebacterium* in dairy, and fungi such as *Rhizopus arrhizus* and *Tausonia pululans*, indicate that these microbes may contribute to development of fermented food environment [90]. Finally, our thorough analyses of microbial communities across a diverse selection of fermented foods allowed us to demonstrate that macronutrient profiles of the raw ingredients of fermented foods are strong determinants of their microbial compositions. The substrate-mediated bacteria-fungi interdependence in our co-occurrence network analysis provides robust evidence that food substrates dictate nutrient availability, imposing strong selection pressures that define the microbial communities and their interactions in traditional fermented foods. Metabolism of nutrients provided by substrates appears to create a highly dynamic ecological process that governs the microbial community formation during food fermentation. The canonical fermenters such as LABs, Bacillus, and yeasts may create favorable conditions for each other’s growth. Yeasts promote the growth of LABs by releasing vitamins and amino acids from foods [91]. Likewise, LAB derived metabolites such as pyruvate, propionate and succinate, promote the proliferation of yeast [92]. Conversely, we also find distinct substrate-specific microbial ecosystems that are not conducive to each other’s growth. Our observations are consistent with inhibitory activities of certain LABs against filamentous fungi, including *Rhizopus*, *Mucor*, and *Penicillium* species [93–95].

Overall, our study identifies a diversity of bacteria and fungi in traditional fermented foods. While some of these bacteria may be involved in food fermentation, others likely contribute to the aesthetics of food. For example, *Brevibacterium* and *Corynebacterium* are associated with appearance [31], *Yarrowia lipolytica* and *Issatchenkia orientalis* enhance flavor [54,76], *Kluyveromyces marxianus* affect aroma [18,79], and *Hannaella oryzae* is linked to the umami taste [21]. Elucidating the functional potential of these microbes may reveal novel fermentation traits pertinent to taste enhancement. In addition, we find several gut-associated bacteria including *Prevotella*, *Ruminococcus, Blautia*, *Faecalibacterium* and *Bifidobacterium* in traditional fermented foods. Interestingly, these are some of the hallmark bacteria that define the gut microbial compositions of non-industrialized populations [96]. This raises an intriguing possibility that traditionally fermented foods may be an important source of gut bacteria for traditional populations. Notably, LABs from dairy have been detected in the gut microbiomes of European populations [97], but whether PUFF microbes can colonize the human gut remains markedly understudied.

Despite these findings, the amplicon sequencing approach we implemented in this study had several limitations that can be mitigated by future metagenomics sequencing. The limited taxonomic resolution offered by amplicon sequencing prohibited us from identifying the source of microbes in fermented foods and whether the same microbial strains are shared across foods originating from different geographies [98]. Although we detected several bacterial and fungal genera, including LABs and yeasts, overlapping across the traditional foods originating from broad geographic regions, we were unable to determine whether these microbes contain strain level variations that may contribute to food fermentation and regional differences. Moreover, some of the microbial genera we detected in our data such as *Klebsiella*, *Citrobacter*, and *Aspergillus* have pathogenic strains. Although presence of pathogenic bacteria in foods that are commonly consumed is unlikely, strain-resolved metagenomes assembled from fermented foods would allow precise functional profiling of bacteria and fungi present in fermented foods [89].

Nevertheless, many of our findings are corroborated by previous metagenomics studies. Several of the PUFF microbes we identified in this study, such as *Klebsiella, Bifidobacteria, Pantoea, Brevibacterium, Corynebacterium, Arthrobacter, Aspergillus,* and *Kluyveromyces* were also identified in previous metagenomic studies [45,85–89]. The substrate specificity of the bacterial composition of fermented foods are also reflected in these metagenomics studies. For example, consistent with our results, *Bacillus* has been reported as the most dominant genera in fermented soy products [85, 89], *Streptococcus* and *Lactobacillus* in fermented dairy [87,89], and *Lactobacillus* and *Leuconostoc* species in fermented vegetables [86,89]. Finally, more precise functional profiling allowed by metagenomics has revealed that plant based fermented foods are enriched for carbohydrate degrading enzymes compared to dairy products [89], which is consistent with our results. While metagenomic profiling of fermented foods is currently at its nascency consisting of small numbers of samples with limited diversity, future studies implementing metagenomics, metabolomics and co-culture experiments are needed to reveal microbes that affect specific aspects of food fermentation [44]. A better understanding of mechanisms underlying interactions between these microbes may allow us to engineer multi-strain microbial communities to improve palatability, preservation, and health benefits.

## Acknowledgements

We express our gratitude to the individuals who contributed their fermented food samples in this study. We thank Nabin Dangol and the Newar community in Khokana, Lalitpur for contributing fermented food samples from Nepal and for facilitating engaging discussions with community members around the importance of this work for preservation of their cultural practices. We are most thankful to Justin L. Sonnenburg for his support throughout this work and for providing helpful critique of the manuscript.

This work was supported by grants from the Bill and Melinda Gates Foundation, New York University Abu Dhabi Startup Fund to ARJ. AG was supported by New York University Abu Dhabi Faculty Research Fund.

## Author Contributions

This study was conceptualized by A.R.J. A.G., B.R., T.W.D, N.A., L.U., collected the samples under the supervision of Y.S.K, N.N.B, and A.R.J. A.G, A.R.A, and A.R.J developed the methods and conducted the analyses. A.G. visualized the data. The first draft of the manuscript was prepared by A.G., R.P., and A.R.J., which was reviewed and edited by all authors.

## Declaration of interests

The authors declare no competing interests.

## Data and code availability

The raw sequencing data for this study will be available from NCBI Sequencing Read Archive (SRA) under submission ID SUB14519890 and BioProject ID PRJNA1126121. The analysis scripts used in this study are available at https://github.com/geneticheritage

## Methods

### Sample Collection

Fermented food samples from Nepal were collected from households, local boutique shops and supermarkets between December 2020 and February 2021. For global references, we obtained ferments of soybean, chili paste, anchovy sauce, cucumber (*oiji*), pickled plum, plum juice, shrimp and *kimchi* from South Korea; a cheese sample was acquired from Kazakhstan; and *Ayib*, *Datta*, *Difo dabo*, *Awaze* and *Injera* samples were retrieved from Ethiopia. Fermented food samples from Ethiopia, Kazakhstan, and South Korea originated from local shops. The samples were stored in sterile 50mL falcon tubes and labelled before shipping to New York University Abu Dhabi (NYUAD). In the NYUAD labs, the fermented foods were aliquoted into multiple 2mL cryogenic tubes and stored at −80°C.

### DNA extractions and ITS/16S amplicon sequencing

DNA extractions were performed using the Zymo PowerSoil DNA Extraction Kit (Manufacture No. D4303). One vial of each sample was suspended in 1mL of Zymo extraction buffer and homogenized in liquid Nitrogen using a mortar and pestle. After each extraction, the mortar was rinsed with a 1mL Zymo extraction buffer and the wash was reserved to detect residual DNA (wash controls). A second set of samples were suspended in the Zymo extraction buffer and vortexed vigorously, before extracting DNA. We included a negative control– extraction reagents without any sample– for every 5 food samples. A detailed protocol for the DNA extractions is available at **[<u>protocols.io</u>]**. In total, we generated 217 samples (homogenized = 64; non-homogenized =90; wash controls = 56; and negative controls = 7) that were shipped on dry ice to Novogene, Singapore, for sequencing. Extracted DNA from each sample was sequenced using multiplexed barcoded primers (515F/806R) for PCR amplification of the 16S rRNA V4 region of bacteria. For the samples with high DNA yield (n=45; homogenized=20; and non-homogenized=25), we performed additional sequencing with ITS1F/ITS2 primers for fungi. Normalized amplicon concentrations were sequenced on a Illumina NextSeq platform to generate 2 × 150 bp paired-end reads. From the 217 samples submitted for sequencing, 13 failed to generate PCR products and all were negative or wash controls. The controls that generated PCR products were sequenced to assess for environmental contamination.

### Computational analysis

The DADA2 16S (1.16) Pipeline Workflow [99] was used to process raw 16S sequences and create a phyloseq object. In order to filter out low quality reads, sequences containing N in the assembled results and/or >2 expected errors were discarded (maxN = 0, maxEE = 2, truncQ = 2), and sequence variants were inferred by pooling reads from all samples (pool = TRUE). All sequences were trimmed from 11 to 150 bp. Forward and reverse reads were then merged to obtain sequence variants. These obtained sequences were filtered to remove chimeric sequences. The Silva database [100] was used for taxonomic classification of Amplicon Sequence Variants (ASV). 11564703 total merged reads and 93,250 ASVs were identified. For quality control, ASVs containing 5 or more reads in more than 10 percent of samples were retained. Furthermore, eukaryotic DNA sequences were removed. A total of 7709741 merged reads passed the quality control, and 861 taxa were identified from a total of 200 samples. Mean (±SD) sequencing depth per sample was 37793 (±17566). The 7 negative and 43 wash controls that generated sequences and passed the quality control were retained. The microbes in the controls were qualitatively different from those of the samples (**Supplementary Figure 1**). Moreover, homogenized and non-homogenized samples showed no significant differences **(Supplementary Figure 2)**. Therefore, controls were removed and homogenized and non-homogenized samples were merged using the merge_samples function from the phyloseq package in R. Then, bacterial ASVs that accounted for less than 0.1% of the total abundances were removed. The final dataset for downstream analyses included 104 bacterial ASVs from 3570678 total reads and 90 samples. Mean (±SD) sequencing depth per sample was 39674(±19200).

In parallel, the DADA2 ITS (1.8) Pipeline Workflow [101] was used to create a phyloseq object from the fungal ITS2 sequences. Since the ITS region is highly variable in length, sequences cannot be trimmed to a fixed length, and primer removal is complicated by the possibility of reads extending into the opposite primer. Therefore, a specialized adapter removal tool, cutadapt [102], was used to filter out primer sequences from the raw reads, and sequences with length greater than 50 were retained (maxN = 0, maxEE = c(2, 2), truncQ = 2, minLen = 50). Sequence variants were inferred by pooling reads from all samples (pool = TRUE), and forward and reverse reads were merged to obtain sequence variants. The obtained sequences were filtered to remove chimeric sequences. The UNITE database [103] was used for taxonomic assignment of the fungal sequences into Amplicon Sequence variants (ASV) respectively. A total of 3475302 merged reads and 2224 taxa were identified from a total of 45 samples. For quality control, ASVs with 5 or more reads in at least 10 percent of samples were retained, and eukaryotic DNA sequences were removed. 2191921 total reads and 193 ASVs passed the quality control. Mean (±SD) sequencing depth per sample was 75583 (±57423). There were no significant differences in microbiome composition of homogenized and non-homogenized fermented food samples **(Supplementary Figure 2)**. As a result, fungal reads from homogenized and non-homogenized samples were merged. Finally, fungal ASVs that accounted for less than 0.1% of the total abundances were removed, and the final dataset contained 91 fungal ASVs from 2139113 total reads and 29 samples. Mean (±SD) sequencing depth per sample was 73763(±56592).

### Bioinformatics and statistical analysis

All statistical analyses were performed in the R environment [R.4.3.0]. Corresponding analysis (CA) was performed on 17 abiotic variables associated with fermented foods using the FactoMineR package [104]. Alpha and beta diversity analyses, and differential abundance of microbial taxa was performed to compare food samples and controls, and homogenized and non-homogenized samples. Alpha diversity was measured using species richness and Shannon’s Diversity Index at the ASV level. Beta diversity was assessed using the weighted UniFrac dissimilarity at the ASV level by log+1 transformation of 16S rRNA gene counts. Differential abundance analysis was done using *differential expression analysis for sequence count data version 2* (DESeq2) package [105]. ASVs with an absolute log2[fold change] > 2 and adjusted p < 0.05 were considered significantly different.

To assess and visualize overall similarity in sample composition, Principal Coordinate Analysis (PCoA) on weighted UniFrac distances were performed for both 16S (n=90 samples) and ITS (n=29 samples) data. To assess whether abiotic factors explained any of the variation observed microbial composition, PERMANOVA tests on bacterial and fungal dissimilarities (with 999 permutations) was performed with 10 metadata variables, including substrate, starter culture, salt, spice and oil usage, fermentation duration, environmental exposure, dryness, shelflife, and country of origin. For ITS data, substrates were grouped into 3 instead of 5 levels for PERMANOVA due to smaller sample size. Plants and cereal were grouped together as plants, and meat and legumes were grouped together. The same grouping was used for subsequent analysis on 16S data with the same 29 samples.

The gap statistic was calculated using the clusGap function from the cluster package with the first four eigenvalues from the PCoA. Clustering based on the ideal number of clusters estimated from the gap statistic was assigned using Partitioning Around Medoids (PAM) clustering. Random forest was employed to determine the precision with which samples could be allocated to clusters assigned through PAM clustering. The performance of the classifier was assessed by generating receiver operating characteristic curves (AUC) using the R-package ROCR [106]. Overall accuracy (true positives+true negatives/false positives+false negatives) of the model was also measured for the test data. Variable Important Factor (VIF) values were calculated for each taxa from the random forest model using the *varImp* function from the *caret* package [107].

Maaslin2 [108] was used to assess differentially abundant bacteria between salted and unsalted foods. Heatmap was generated for ASVs with adjusted p < 0.05 and coef>2. T-tests and ANOVA were used to compare means across groups when assumptions of normality were met (p > 0.05 using Shapiro test), while Mann-Whitney and Kruskal-Wallis tests were utilized when normality assumptions were violated. Fisher’s test was used to assess enrichment of substrates in PCoA clusters. Brown Forsythe test was used to estimate the equality of group variances.

### Functional Analysis

Picrust2 was used to predict functional pathways using 16S sequences. The resulting KO terms were filtered for those that accounted for more than 0.05% of the total pathway abundance. KO terms were converted to KEGG pathways and Linear Models for Differential Analysis (LinDA) was used to calculate the differentially abundant KEGG pathways using the ggpircust2 package [109]. Differential abundance analysis was performed using 3 substrate levels: plants (cereals and vegetables); dairy; and legumes, meat, and seafood. Plants were used as reference, which generated differentially abundant functions between plants and legumes/meat/seafood, and plants and dairy. The reference was changed to dairy to calculate differences between legumes/meat/seafood and dairy. Multiple testing corrections were performed by computing FDRs using the Benjamini–Hochberg method. Functions with FDR corrected p <0.05 were considered significantly different.

### Co-occurrence networks

FastSpar [110] was used to estimate a correlation network on ASVs and calculate p values. Parameters included 1000 bootstrap correlations (--number 1000) and 1000 permutations for p value estimates (-- permutations 1000). Correlations were filtered for ≤0.3 or ≥0.3 and permuted p-values ≤0.05 were considered significant. A correlation heatmap was generated using the pheatmap package. Taxa with at least one significant correlation was considered to be part of co-abundant groups (CAGs). Clusters of taxa forming CAGs were identified using hierarchical clustering, and the ideal number of CAGs were estimated using gap statistics on hierarchical clustering. Network plots were generated using igraph (111). Focal taxa was assessed by measuring the strength (number of edges weighted by their correlation), betweenness (the number of shortest paths that go through a given bacterial ASV), and eigenvector centrality, which measures the node’s importance based on its connections to other highly connected nodes in the network **(Supplementary Figure 8).** CAG properties were assessed by estimating connectedness and transitivity. Connectedness was calculated as the number of edges of each node (ASV) within a CAG, normalized by the total number of nodes in the CAG. Transitivity was estimated, for each node, as the proportion of their neighbors connected to each other. Both metrics evaluate how densely taxa are interconnected within each CAG. These measures were calculated using respective functions from the igraph package, with support from Shizuka’s network analysis tutorial github page [112].

### Identification of candidate species

We used the SILVA database to generate genus level identifications, and EZ BioCloud and NCBI 16S rRNA databases to generate species level identifications for the bacterial ASVs. The 16S rRNA gene sequences from fermented food samples were queried against the EZBioCloud Database of reference 16S rRNA gene sequences (version 2023.08.23). Species level assignments were made for sequences with ≥99.3% unambiguous match to a type (T) sequence. Likewise, the sequences were also queried against the NCBI database using the BLAST tool. Species level assignments were made for sequences with 100% unambiguous match with a maximum of 2 subject sequences. Out of 104 sequences, 52 were unable to be resolved at the species level using either databases. 34 sequences exhibited matching results for the same species in both BLAST and EZ BioCloud Database, consequently leading to their resolution at the species level. BLAST successfully characterized 16 sequences that remained uncharacterized by the EZ BioCloud Database. In this case, species-level identification was done using BLAST. For 2 sequences, EZ BioCloud and BLAST resulted in conflicting species level identifications. In this case, both the species recommendations were used **(Supplementary Table 4)**. The UNITE database was used to generate species level identifications for the fungal ASVs.

## Supplementary Figures

**Supplementary Figure 1. There are no significant differences between homogenized and non-homogenized samples. (A)** DNA concentrations were higher for food samples compared to controls (p<0.05, *ANOVA*). Out of 217 samples, 204 generated bacterial 16S sequences. The ones that failed to generate sequences were all controls. Only 45 samples yielded enough DNA (>10 ng/uL) to generate fungal ITS reads. **(B)** Principal Coordinate Analysis (PCoA) of Weighted UniFrac dissimilarity revealed controls deviated from the fermented food samples (PERMANOVA, p= 0.0001). Alpha diversity between controls and fermented foods measured using Shannon Diversity Index **(C)** and Species Richness **(D)** showed significant differences between control and test groups (Kruskal Wallis test at rarefaction depth =10000, p =0.0001 for both). **(E)** Differential abundance analysis between the controls and ferments using DESeq2 found 614 of 861 bacterial ASVs were differentially abundant in fermented food samples compared to controls (FDR adjusted p value <0.05). ASVs belonging to orders Lactobacillales (LABs) and Bacillales, the canonical bacteria implicated in food fermentation, were more abundant in the food samples compared to controls. These results underscore the integrity of microbial communities curated from the ferments. **(F)** Principal Coordinate Analysis (PCoA) of the food samples only using weighted UniFrac distances from 16S rRNA reads revealed no significant differences between the 64 homogenized food samples and their non-homogenized replicates. **(G)** Bacterial alpha diversity was also comparable (p<0.05, wilcox test) between homogenized and non-homogenized samples. **(H)** None of the bacterial ASVs showed significant differences in their relative abundances between the homogenized and non-homogenized samples (FDR adjusted p>0.05, DESeq2). **(I)** Principal Coordinate Analysis (PCoA) performed using weighted UniFrac distances from ITS2 reads also revealed no significant differences between the 16 homogenized food samples and their non-homogenized replicates. **(J)** Fungal alpha diversity was not significantly different (p<0.05, wilcox test) between the homogenized and non-homogenized samples. **(K)** The relative abundances of none of the fungal ASVs were significantly different between the homogenized and non-homogenized samples (FDR adjusted p>0.05, DESeq2).

**Supplementary Figure 2. Traditional fermented foods exhibit heterogeneity in their microbial composition. (A)** Distribution of the number of ASVs belonging to Canonical Fermenters in each food sample. **(B)** Relative abundances of Canonical Fermenters across the five substrate types. **(C,D)** Relative abundance of key PUFF bacteria and fungi across fermented food substrates.

**Supplementary Figure 3. Factors associated with bacterial composition in traditionally fermented foods.** Principle Coordinate Analysis (PCoA) of weighted unifrac dissimilarity matrix generated from 16S rRNA reads from 90 traditional fermented foods considering all 104 bacterial ASVs **(A)** and canonical fermenters only **(B)**. Colors are mapped to country of origin and shapes are mapped to substrate type. **(C-D)** Gap statistics from the partition around medoids (PAM) clustering performed using the top four PCo axes obtained using all 104 ASVs **(C)** and canonical fermenters only **(D)**. **(E)** Effect sizes (R^2^) of factors (ingredients and preparation methods) and the bacterial communities in traditional fermented foods obtained from PERMANOVA on weighted dissimilarity matrix generated from 16S reads from 90 traditional fermented foods considering all taxa (red), and canonical fermenters only (blue). Asterisks (*) indicate factors that are significantly associated with the bacterial community (p <0.05). The effect of geography, which is significant when all 104 microbes are included in the model (p=0.0001, *PERMANOVA*) becomes negligible when only canonical fermenters are included (p>0.05, *PERMANOVA*). **(F)** AUC curves from random forest classifiers used to distinguish the 4 bacterial clusters obtained from PAM clustering. **(G)** The relative abundances of 20 most important bacterial ASVs used by the random forest model to distinguish samples into 4 clusters across substrate types.

**Supplementary Figure 4. Substrate plays a less prominent role in the establishment of fungal community in traditional fermented foods. (A-B)** Principle Coordinate Analysis (PCoA) of weighted unifrac dissimilarity matrix generated from ITS2 reads from 29 traditional fermented foods considering all 91 fungal ASVs **(A)** and canonical fermenters (Saccharomycetales) only **(B)**. Circles represent samples color coded by substrate types. **(C-D)** Gap statistics from the partition around medoids (PAM) clustering performed using the top four PCo axes obtained using all 91 ASVs **(C)** and canonical fermenters only **(D)**. **(E)** AUC curves from random forest classifiers used to distinguish the 3 bacterial clusters obtained from PAM clustering for A. **(F)** Effect sizes (R^2^) of factors (ingredients and preparation methods) and the fungal communities in traditional fermented foods obtained from PERMANOVA on weighted dissimilarity matrix generated from 16S reads from the 29 traditional fermented foods considering all taxa (red), and canonical fermenters only (blue). Asterisks (*) indicate factors that are significantly associated with the fungal community (p <0.05). **(G)** The relative abundances of 20 most important fungal ASVs used by the random forest model to distinguish samples into the 3 clusters across substrate types.

**Supplementary Figure 5. Bacterial composition of the same 29 samples for which the fungal data is available. (A-B)** Principle Coordinate Analysis (PCoA) of weighted unifrac dissimilarity matrix generated from 16S rRNA reads from 29 traditional fermented foods which have both 16S rRNA and ITS2 data and considering all taxa **(A)** and considering canonical fermenters only **(B)**. **(C)** Effect sizes (R2) of factors that were included in PERMANOVA analysis on weighted dissimilarity matrix generated from 16S reads from 29 traditional fermented foods which have both bacterial and fungal data, considering all taxa (red), and canonical fermenters only (blue). Factors with a statistically significant association with the dissimilarity matrix (p <0.05) are marked with an asterisk (*).

**Supplementary Figure 6. Co-occurrence analysis reveals densely connected bacterial and fungal networks. (A)** A heatmap showing co-occurrence patterns among the 104 bacterial taxa. Colors indicate strength of co-occurrences. **(B)** Network plot showing significantly positive associations (R^2^>0.3, p<0.05) among bacterial ASVs. **(C)** Co-occurrence patterns among the 91 fungal ASVs. **(D)** Network plot showing significantly positive associations (R^2^>0.3, p<0.05) among the fungal ASVs.

**Supplementary Figure 7. Interconnectedness between fermented food microbes. (A)** Bacterial CAGs are significantly different in their connectedness (ANOVA<2e-16; all CAGs are significantly different in pairwise comparisons except those labeled “ns” (p<0.05, Tukey HSD). Connectedness is measured as the number of significant associations an ASV has with other ASVs in its CAG normalized by the total number of ASVs in the CAG. **(B)** Bacterial CAGs are significantly different in their transitivity (ANOVA<2e-16; significantly different CAGs in pairwise comparisons are marked with an *, p<0.05, Tukey HSD test). For each CAG, transitivity measures the proportion of ASVs connected with their neighbors that are connected to each other. **(C)** Fungal CAGs show significant differences in their connectedness (ANOVA<2e-16). Two fungal CAGs are significantly different in pairwise comparisons (p>0.05, Tukey HSD) **(D)** Fungal CAGs are significantly different in their transitivity (ANOVA<2e-16). Significantly different CAGs in pairwise comparisons are marked with an *, p<0.05, Tukey HSD test).

**Supplementary Figure 8. Identification of focal species of bacteria and fungi in traditional fermented foods.** Identification of focal microbes within each CAG by assessing three measures of centrality, namely strength, betweenness, and eigenvector centrality for ASVs belonging to bacteria (**A-E**) and fungi (**F-J**). Size of the circles represent normalized values for each of these measures and ranges from 0 to 1. Text inside circles represents actual strength, betweenness, and eigenvector centrality values of bacterial and fungal ASVs within their respective CAGs.

## Supplementary tables

**Supplementary Table 1.** The description of samples, substrate, preparation methods and macronutrient profiles obtained from current literature.

**Supplementary Table 2:** Microbial putative pathways that are differentially abundant among substrates (plants as reference).

**Supplementary Table 3 :**Microbial putative pathways that are differentially abundant among substrates (dairy as reference).

**Supplementary Table 4:** Species level identification for bacterial ASVs using EZ BioCloud and BLAST.

## Notes

### Competing Interest Statement

The authors have declared no competing interest.

