## Supplementary figures and images for "Ecological factors that drive microbial communities in culturally diverse fermented foods"

### Supplementary Figure 1

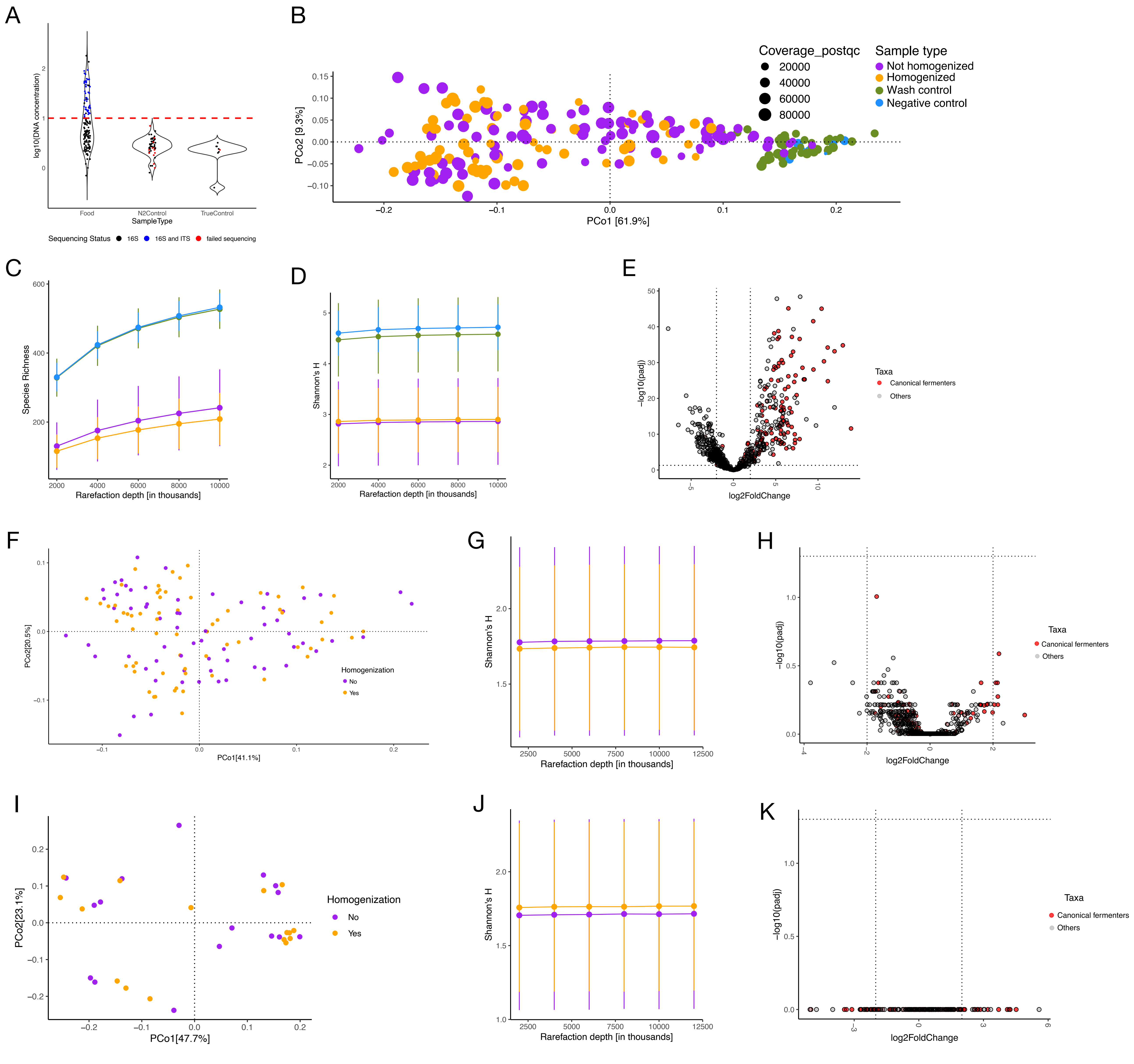

### Supplementary Figure 2

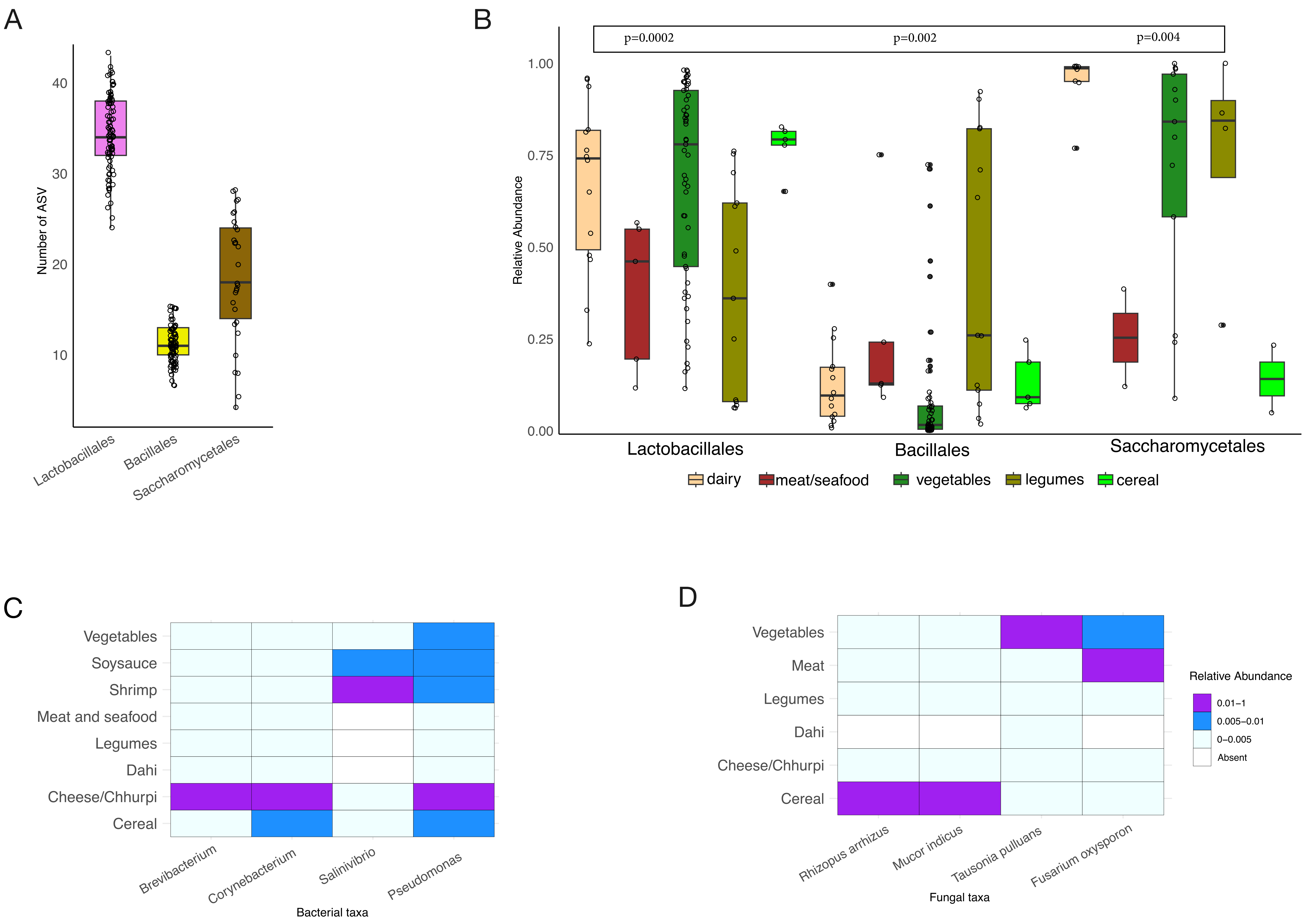

### Supplementary Figure 3

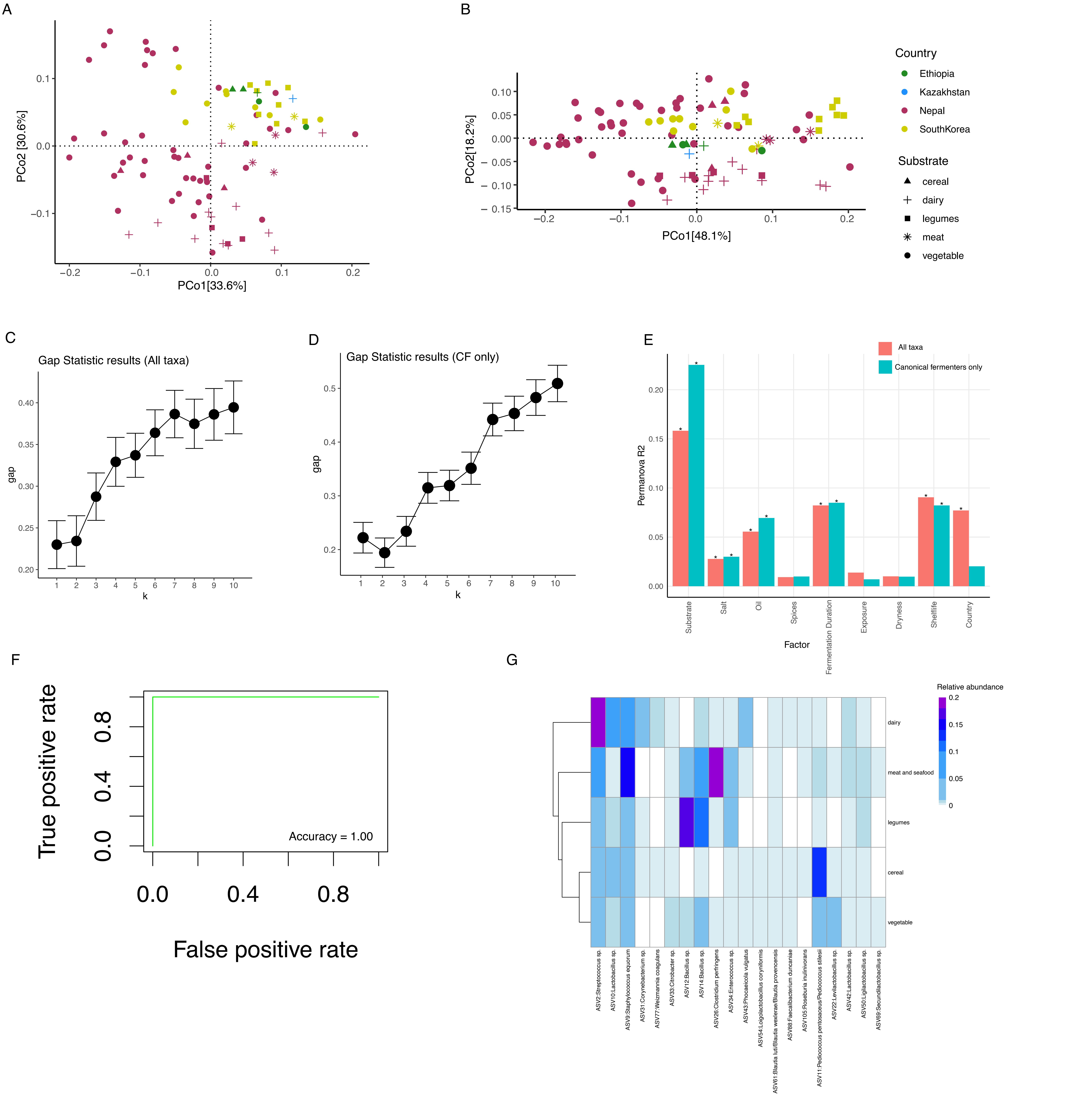

### Supplementary Figure 4

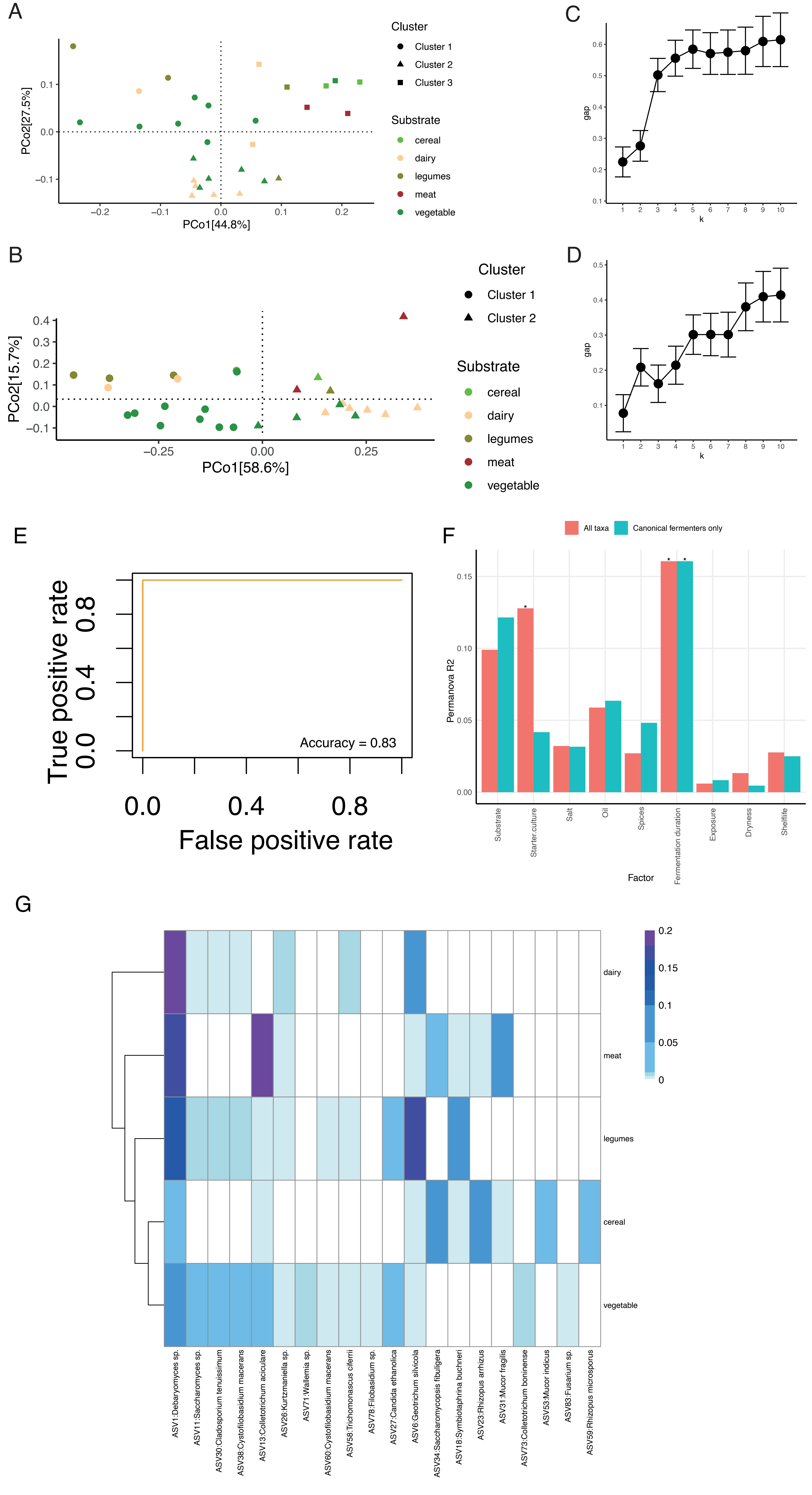

### Supplementary Figure 5

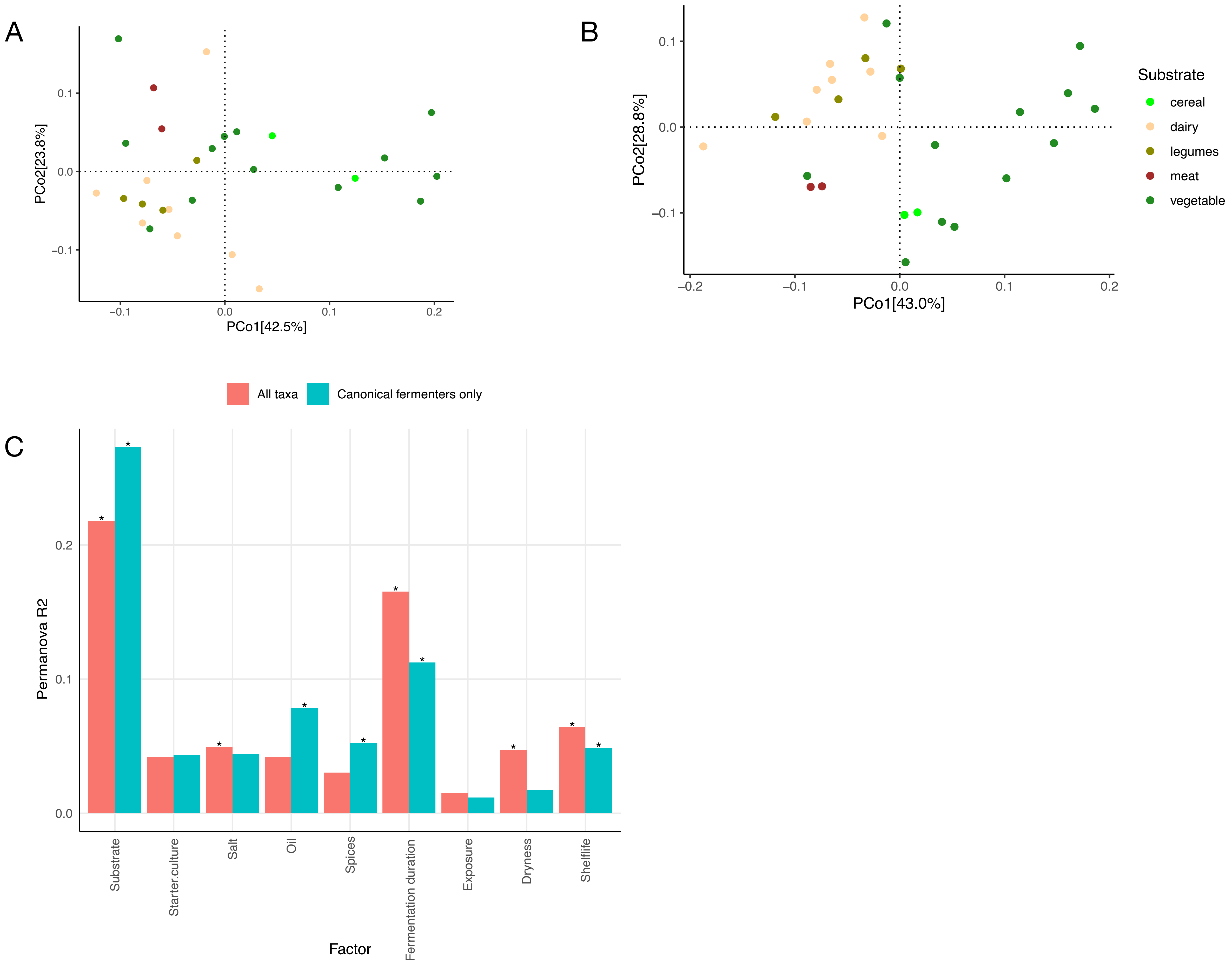

### Supplementary Figure 6

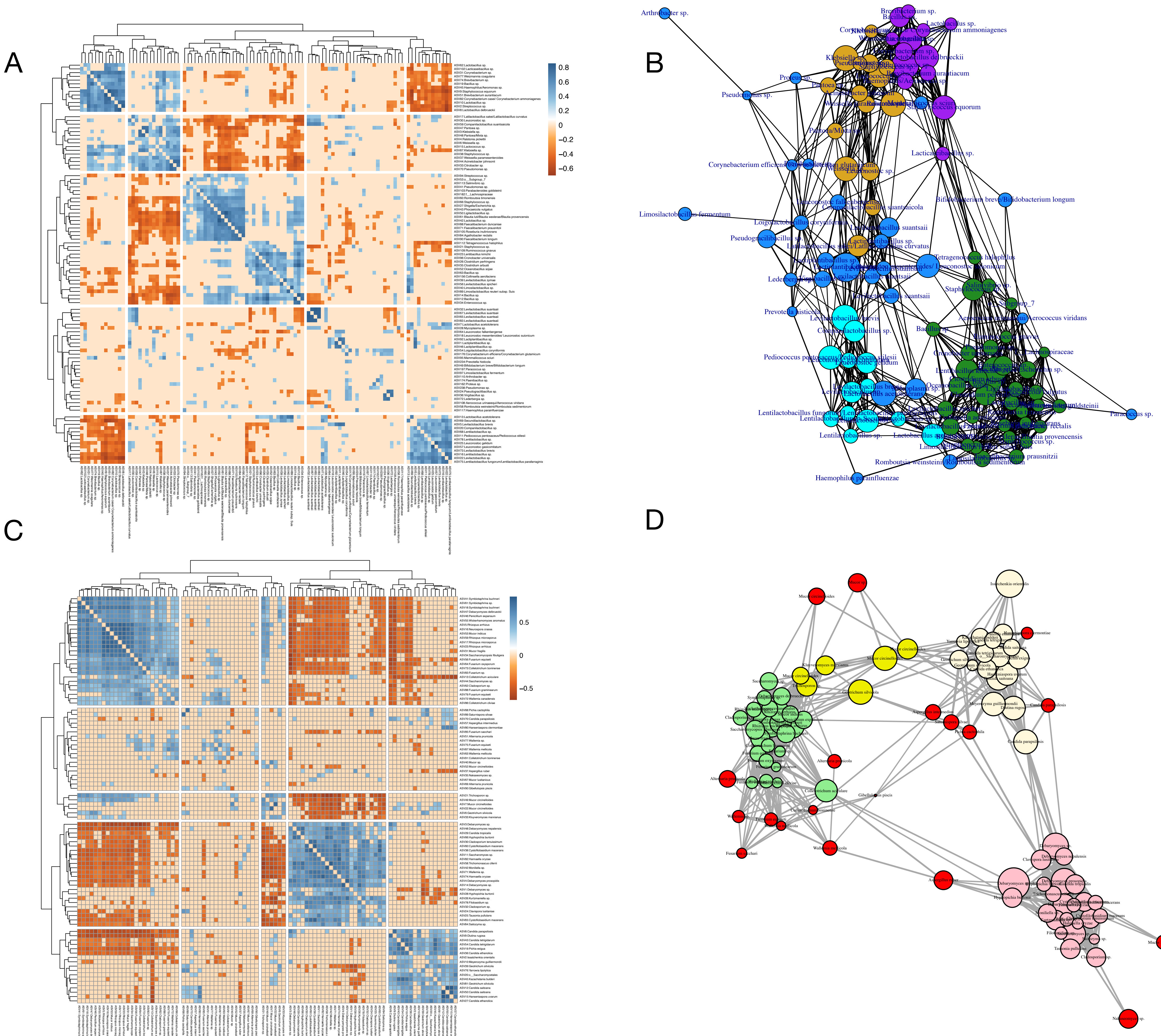

### Supplementary Figure 7

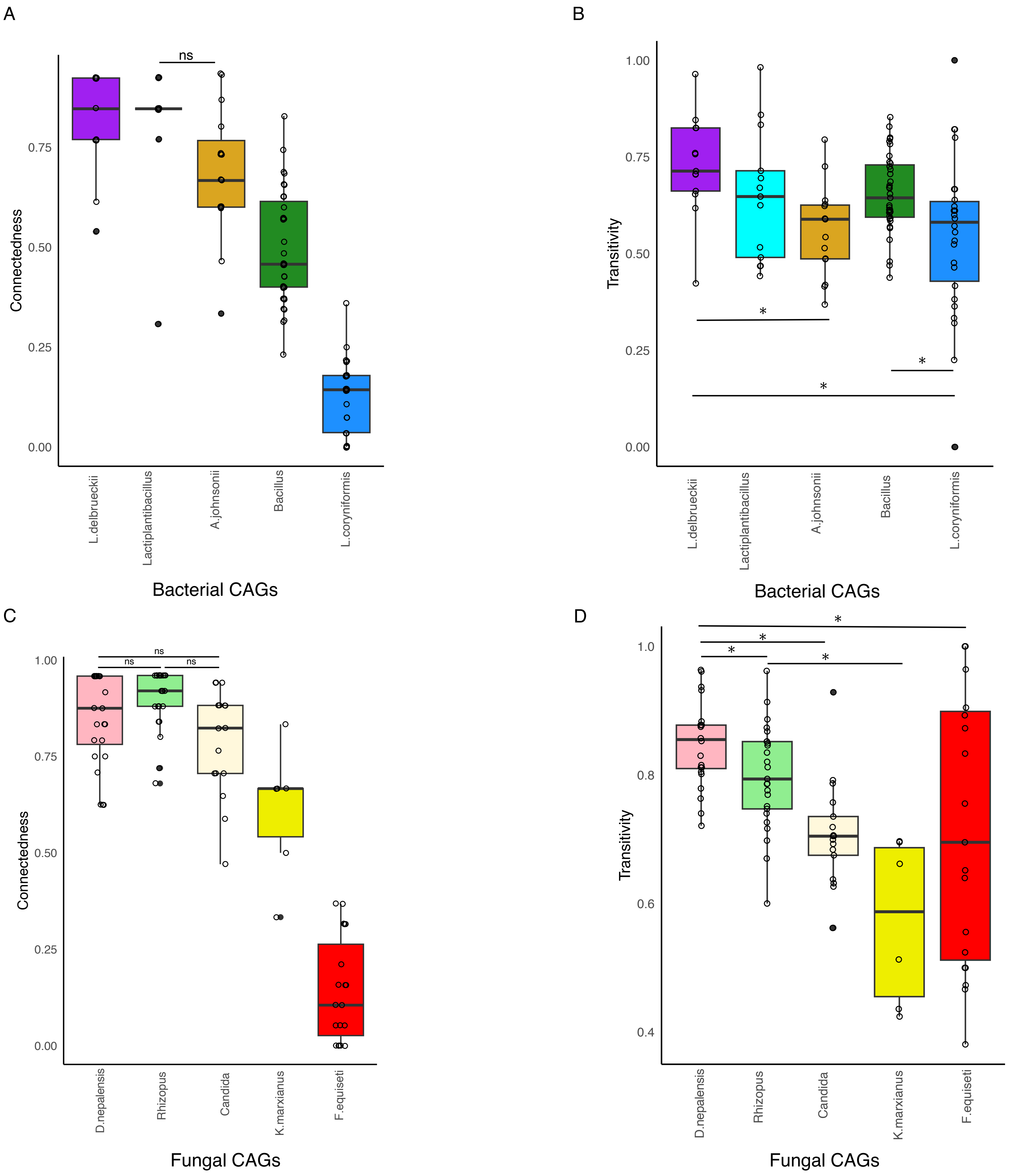

### Supplementary Figure 8

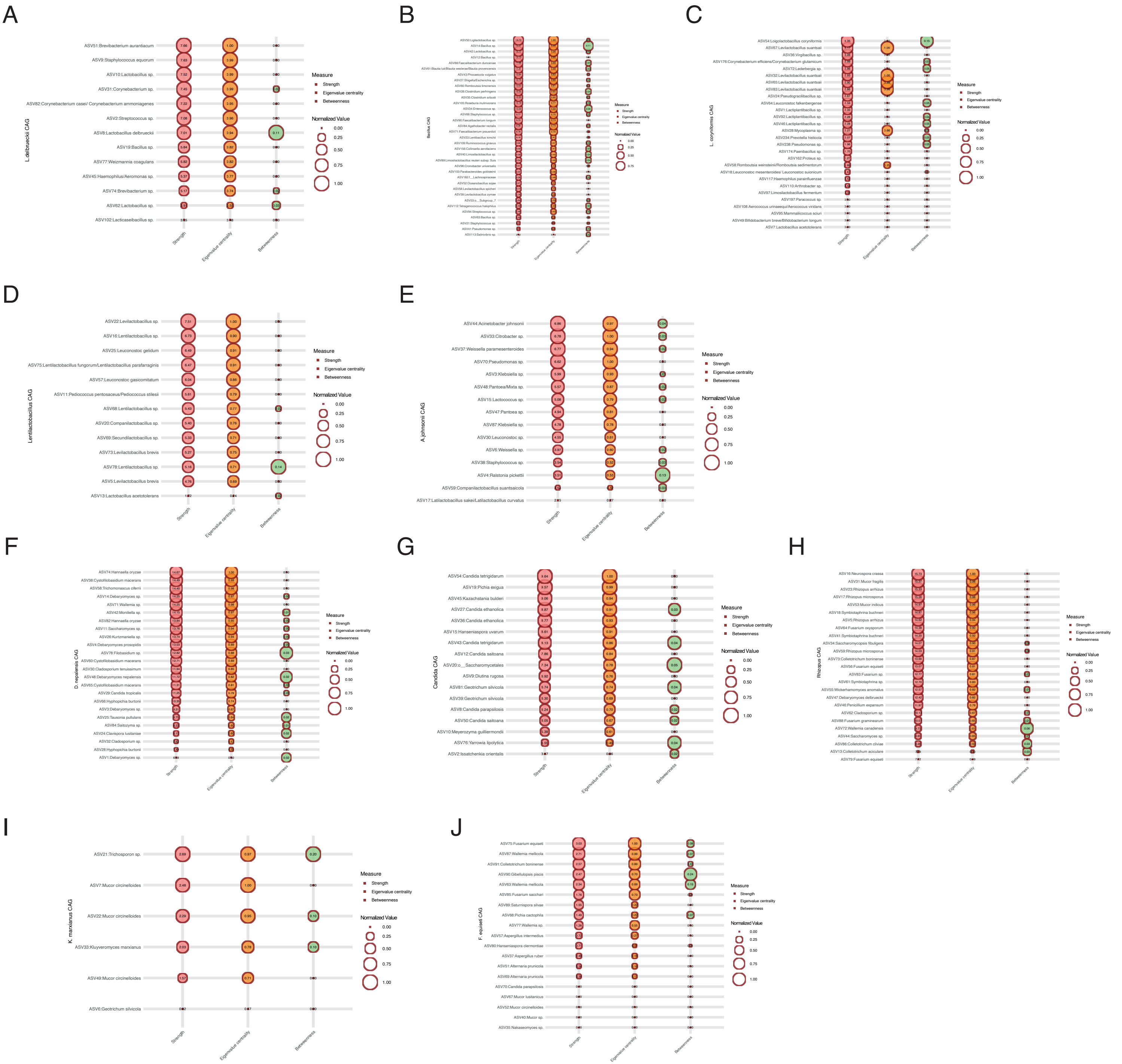
